# IL-17A both initiates (via IFNγ suppression) and limits the pulmonary type-2 immune response to nematode infection

**DOI:** 10.1101/827899

**Authors:** Jesuthas Ajendra, Alistair L. Chenery, James E. Parkinson, Brian H. K. Chan, Stella Pearson, Stefano A. P. Colombo, Louis Boon, Richard K. Grencis, Tara E. Sutherland, Judith E. Allen

## Abstract

*Nippostrongylus brasiliensis* is a well-defined model of type-2 immunity but the early lung-migrating phase is dominated by innate IL-17A production and neutrophilia. Using *N. brasiliensis* infection we confirm previous observations that *Il17a*-KO mice exhibit an impaired type-2 immune response. Neutrophil depletion and reconstitution studies demonstrated that neutrophils contribute to the subsequent eosinophilia but are not responsible for the ability of IL-17A to promote type-2 cytokine responses. Transcriptional profiling of the lung on day 2 of *N. brasiliensis* infection revealed an increased *Ifnγ* signature in the *Il17a*-KO mice confirmed by enhanced IFNγ protein production. Depletion of early IFNγ rescued type-2 immune responses in the *Il17a*-KO mice demonstrating that IL-17A-mediated suppression of IFNγ promotes type-2 immunity. Notably, when IL-17A was blocked later in infection, the type-2 response increased. IL-17A regulation of type-2 immunity was lung-specific and infection with *Trichuris muris,* revealed that IL-17A promotes a type-2 immune response in the lung even when a parasite lifecycle is restricted to the intestine. Together our data reveal IL-17A as a major regulator of pulmonary type-2 immunity which supports the development of a protective type-2 immune response but subsequently limits the magnitude of that response.

## INTRODUCTION

Innate and adaptive sources of interleukin-17A (IL-17A) are responsible for a range of neutrophil-associated inflammatory conditions as well as protection from many bacterial and fungal pathogens^1,2^. In contrast, type-2 immunity is required for effective control of most helminth infections^3^ and is characterized by eosinophilic inflammation and the cytokines IL-4, IL-5 and IL-13. When both type-2 and IL-17 responses are present during helminth infection enhanced pathology is observed, as shown for human schistosomiasis^4,5^ and onchocerciasis^6^. The detrimental relationship between IL-17A and type-2 associated diseases has also been extensively documented in allergic asthma in which the most severe symptoms occur in patients with both high Th2 and Th17 cell responses^7^. Critically, type-2 cytokines can actively suppress IL-17A production which may be an important feedback mechanism to avoid extreme IL-17A-driven pathology^8–10^. Despite evidence for an important relationship between IL-17A and type-2 immune responses during chronic disease, how these responses are connected remains poorly understood.

We and others have demonstrated a prominent role for IL-17A during infection with the lung-migrating nematode *Nippostronglyus brasiliensis*^8,11^, a well-defined pulmonary model of type-2 immunity. After entering the host via the skin, *N. brasiliensis* larvae migrate through the lung, causing tissue damage and haemorrhage. IL-17RA-dependent neutrophil recruitment is largely responsible for the lung damage in this model^11^. We previously found that the chitinase-like protein Ym1 induces expansion of IL-17A-producing γδ T cells and Ym1 blockade or IL-17A-deficiency protects mice from peak lung damage^8^. More surprising was our finding that Ym1 neutralisation or IL-17A-deficiency prevents the development of a full type-2 response during *N. brasiliensis* infection^8^.

The notion that IL-17A is required for development of a type-2 response appears counter to the evidence that type-2 cytokines suppress IL-17A production^9,10^. However, previous studies using murine models of allergic inflammation also show impaired type-2 immunity in the face of IL-17A-deficiency^12,13^ or blockade^14^. In an infection or injury context, the specific tissue as well as timing might all play decisive roles in whether IL-17A augments or suppresses type-2 responses. We therefore used *N. brasiliensis* infection to address the contribution of γδ T cell-derived IL-17A and neutrophils to the development of a subsequent type-2 immune response in the lung. We found that IL-17A suppressed early IFNγ production and that this suppression was essential for the optimal development of a type-2 response. Although neutrophils acted as a major driver of subsequent lung eosinophilia, neutrophils were not responsible for IL-17A-mediated enhancement of type-2 immunity. Once the type-2 response was established, IL-17A acted as a negative regulator, revealing distinct roles during innate and adaptive stages of the response. Notably, *Trichuris muris,* a nematode restricted to the gastro-intestinal tract also induced a lung type-2 response that was IL-17A-dependent. However, we found no evidence that IL-17A regulated the intestinal type-2 response. Thus, IL-17A serves as a lung-specific regulator of the type-2 immune response.

## RESULTS

### IL-17A-deficient mice mount a diminished type-2 response

In keeping with the known ability of *N. brasiliensis* to induce a strong pulmonary type-2 immune response on day 6 post infection (d6pi), we found the BAL and lungs of C57Bl/6 mice to be dominated by eosinophils (Suppl. Fig. 1A, B). This response was accompanied by elevated numbers of CD4^+^ T cells as well as induction of Group 2 Innate lymphoid cells (ILC2, Suppl. Fig. 1C, D). The establishment of a type-2 response was further confirmed by increased type-2 cytokine expression by CD4^+^ T cells and gene expression in whole lung (Suppl. Fig. 1E, F). As we and others previously reported^8,11^, infected mice exhibited increased IL-17A production within the first 48 h post infection (Fig. 1A) and consistent with previous reports, the main source of IL-17A was γδ T cells^8^. On d2pi the BAL consisted mainly of neutrophils (Suppl. Fig.1A), which, together with *N. brasiliensis* larvae migration, is known to cause acute lung injury^11^.

**Figure 1:**
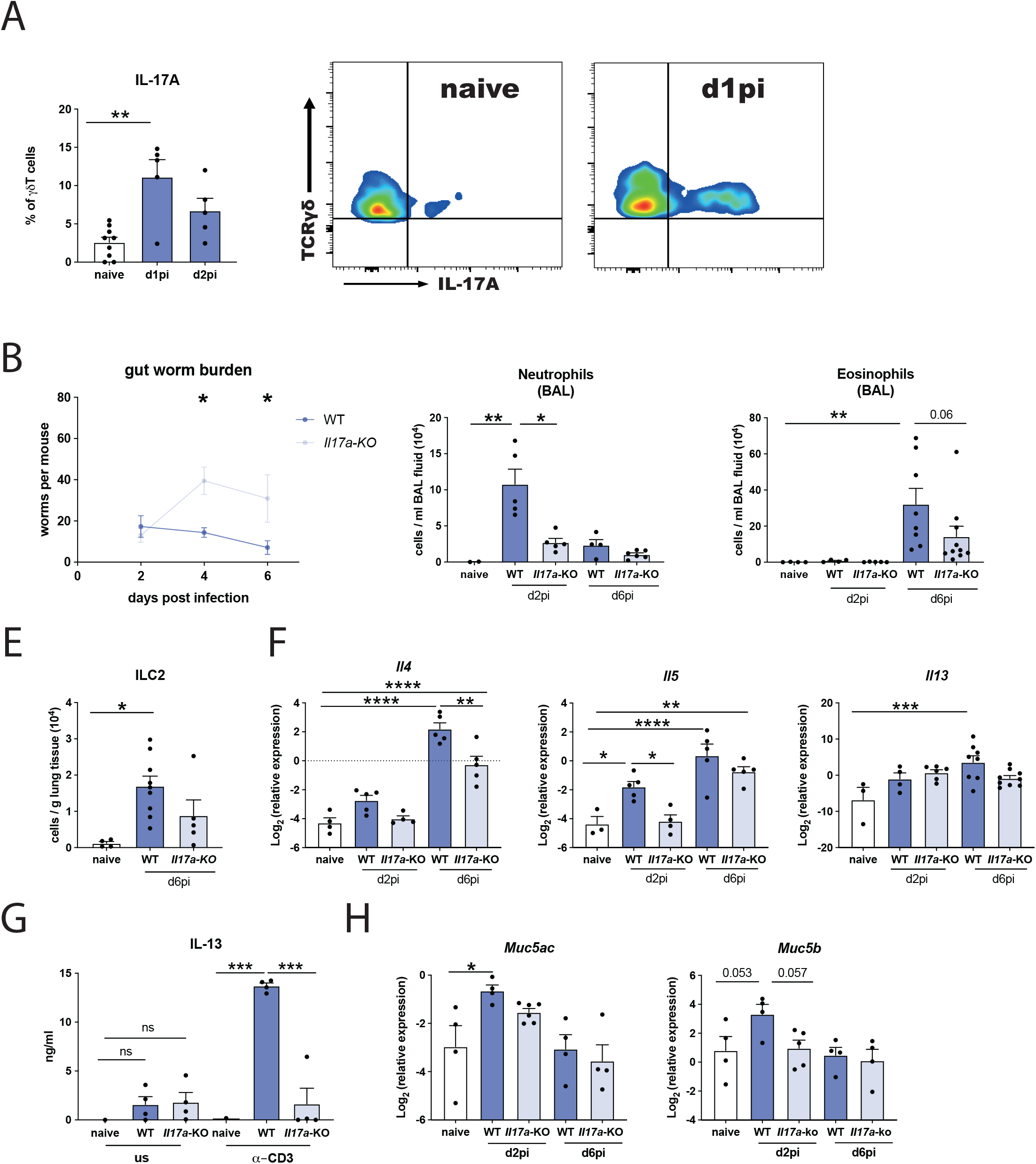
Mice deficient in IL-17A mount a diminished type-2 response at the site of infection. C57BL/6J (WT) and *Il17a-*KO mice were infected with 250 *N. brasiliensis* L3s and cell frequencies and cytokines were measured at different time points post infection compared to WT naïve mice. Frequencies of IL-17A-producing γδ T cells on d1pi and d2pi and representative flow-plot at d1pi (**A**). Worm burden in small intestine assessed in WT and *Il17a-* KO mice on days 2, 4 and 6 post *N. brasiliensis* infection (**B**). Absolute count of neutrophils (Ly6G^+^CD11b^+^) (**C**), eosinophils (SiglecF^+^CD11b^+^ Cd11c^−^) in bronchoalveolar lavage (BAL) **(D**) and lung ILC2 (Lineage^−^ KLRG^+^CD127^+^CD90.2^+^ST2^+^) as measured via flow cytometry (**E**). Relative mRNA expression of cytokines *Il4*, *Il5* and *Il13* in whole lung as quantified by qRT-PCR (log2 expression relative to *actb* (β-actin)) (**F**). Secreted IL-13 levels from unstimulated or 72h α-CD3 treated single-suspension lung cells (**G**). Relative mRNA expression of the mucin genes *Muc5ac* and *Muc5b* in whole lung (log2 expression relative to *actb*) (**G**). Data are representative (mean ± s.e.m.) of at least 3 individual experiments (**A**, **C**, **E**-**F**) or pooled of two experiments (**D**, **G**, **H**) with at least 3 mice per group or pooled data from three experiments (**B**). Data were tested for normality using Shapiro-Wilk test and analysed using one-way ANOVA followed by Sidak’s multiple comparisons test for selected groups. NS – not significant. Data in (**F, H**) were log2 transformed to achieve normal distribution and statistical tests were performed on transformed data \**P*<0.05, \*\**P*<0.01, \*\*\**P*<0.001, \*\*\*\**P*<0.0001.

To investigate the role of IL-17A during the development of type-2 immune responses, we infected *Il17a-*KO mice and WT C57BL/6J controls with *N. brasiliensis* L3’s. Larvae leave the lung within 48 h and are expelled from the gut within 6-8 days. Consistent with our previous findings^8^ *Il17a-*KO mice were significantly more susceptible to infection exhibiting an intestinal worm burden almost twice as high as in the WT controls on d4pi and d6pi (Fig. 1B). As expected, the early d2 neutrophilia in response to *N. brasiliensis* infection was muted in the *Il17a-*KO mice relative to the WT controls (Fig. 1C). Between d2pi and d6pi, there was a switch from neutrophilic to eosinophilic responses in the lungs (Suppl. Fig 1A). Whilst increased eosinophil numbers were observed in all infected d6pi mice relative to naïve animals, this increase was less evident for *Il17a-*KO mice (Fig. 1D). ILC2 displayed a similar pattern, as cell numbers significantly increased with infection in WT but not in *Il17a-KO* mice (Fig. 1E). Additionally, WT mice exhibited an increased type-2 cytokine expression signature at d6pi, whilst *Il17a-*KO mice had reduced *Il4* expression, significantly delayed increase in *Il5* expression and muted *Il13* expression (Fig. 1F). IL-13 protein levels in whole lung restimulated with α-CD3 on d6pi were significantly higher in WT mice compared to naïve controls whilst *Il17a-*KO mice completely failed to upregulate IL-13 levels (Fig. 1G). We also measured expression levels of the major mucins in the lung because host mucin production is another feature of protective type-2 responses^15^. *N. brasiliensis* infection drove an early increase in mucins *Muc5ac* and *Muc5b* expression in the lungs of WT mice at d2pi corresponding to a timepoint when the larvae are transitioning from the lungs. Expression of *Muc5ac* was significantly increased in WT mice compared to naïve controls, but reduced in *Il17a-*KO mice, although not significantly (Fig. 1H). A similar pattern was observed for *Muc5b*, which was upregulated in WT mice by d2pi but failed to increase in *Il17a*-KO mice (Fig. 1H). By d6pi, when the worms had already migrated to the small intestines, increased mucin expression was no longer observed in infected animals (Fig. 1H). Together these data demonstrate an impairment of the pulmonary type-2 immune response during helminth infection in the absence of IL-17A.

### IL-17A regulates T cell activation and polarization during *N. brasiliensis* infection

Next we aimed to determine whether the impact of IL-17A-deficiency on type-2 immunity was due to changes in T cell activation or polarisation during *N. brasiliensis* infection. Using flow cytometry, we observed a reduction in total numbers of CD4^+^ T cells on d7pi in the lungs of *Il17a-*KO mice compared to WT controls (Fig. 2A). In contrast, there were no significant differences in CD4^+^ T cell numbers in the lung-draining lymph nodes (Fig. 2A). We also examined expression of the Th2 transcription factor GATA3. While WT mice showed a significant increase in frequency and absolute numbers of GATA3^+^CD4^+^ T cells upon infection, *Il17a-*KO mice had a significantly lower frequency of GATA3^+^CD4^+^ T cells and failed to upregulate these cells on d7pi compared to WT controls (Fig. 2B).

**Figure 2:**
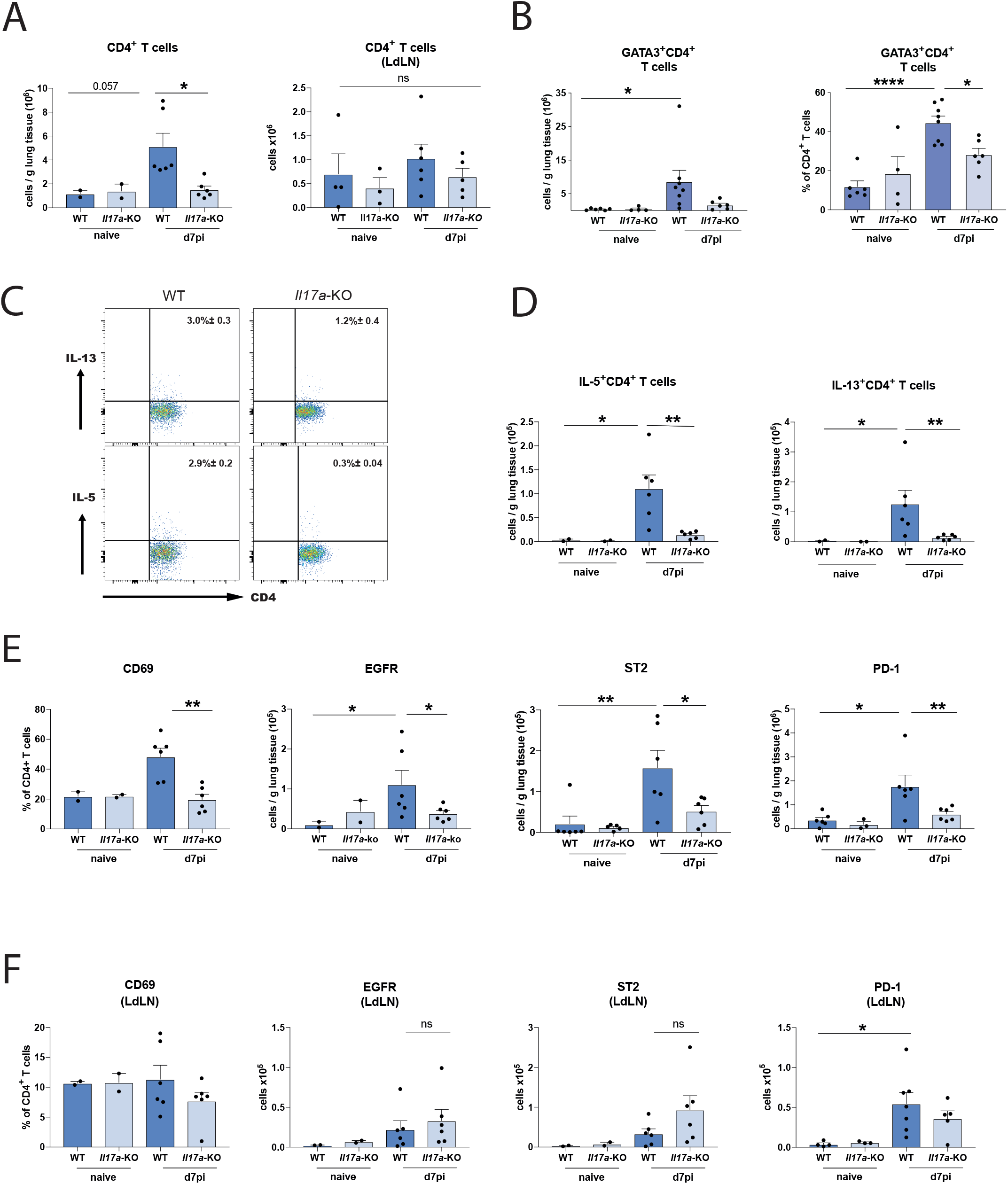
IL-17A regulates T cell activation and polarization during *N. brasiliensis* infection. C57BL/6J (WT) and *Il17a-KO* mice were infected with 250 *N. brasiliensis* L3 larvae or left uninfected (naïve) and live, single CD4^+^ T cells were phenotyped using flow cytometry at d7pi in lung and lung draining lymph nodes (LdLN). Absolute numbers of live CD4^+^ T cells in lung tissue and LdLN (**A**). Frequency and absolute numbers of GATA3^+^ CD4^+^ T cells in the lung (**B**). Representative flow-plots showing the frequency of IL-5 and IL-13 production by CD4^+^ T cells d7pi in lung from WT and *Il17a-KO* infected mice (**C**). Absolute numbers of IL-5^+^ and IL-13^+^ CD4^+^ T cells in the lung (**D**). Expression of CD69 on CD4^+^T cells and absolute numbers of EGFR^+^CD4^+^T cells, ST2^+^CD4^+^T cells and PD-1^+^CD4^+^T cells in lung (**E**) and LdLNs (**F**). Data are representative (mean ± s.e.m.) of at least 3 individual experiments with at least n=2 mice in naive groups and n=4 mice in infected groups (per experiment). Data was tested for normality using Shapiro-Wilk test and analysed using one-way ANOVA followed by Sidak’s multiple comparisons test for selected groups. NS not significant, \**P*<0.05, \*\*\*\**P*<0.01.

Not only were there fewer GATA3^+^ CD4^+^ T cells in the lungs of *Il17a-*KO mice, the CD4^+^ T cells of *Il17a*-KO mice were producing significantly less IL-13 and IL-5 (Fig. 2C). Strikingly, by d7pi, IL-5^+^ and IL-13^+^ CD4^+^ T cell numbers failed to increase in response to infection in *Il17a*-KO mice (Fig. 2D). At the same time point post-infection, we found that expression of the activation marker CD69 was upregulated on CD4^+^ T cells in the lungs of WT but not *Il17a-*KO mice (Fig. 2E). However, CD69 did not differ between all tested groups in the lung-draining lymph nodes (Fig. 2F). Recently, Minutti et al. showed that epidermal growth factor receptor (EGFR) in complex with ST2 on T cells allows for IL-33-induced IL-13 production at the site of *N. brasiliensis* infection^16^. We therefore analysed surface expression of EGFR and ST2 on lung and lung-draining lymph node T cells. *N. brasiliensis* infection increased the number of CD4^+^ T cells expressing these markers in the lung, but this increase was significantly reduced in *Il17a-*KO mice. (Fig. 2E). Changes to ST2 and EGFR expression between WT and KO mice were not observed in the lung-draining-lymph nodes (Fig. 2F). We also measured PD-1 expression, an important regulator of T cell function during helminth infection^17,18^. *N. brasiliensis* infection in WT mice led to increased numbers of CD4^+^ T cells expressing PD-1 in the lung with significantly fewer of these cells in *Il17a-*KO mice (Fig. 2E). Whilst there were also increased numbers of PD-1^+^CD4^+^ T cells in the lymph node following infection of WT mice, these were not altered by IL-17A deficiency (Fig. 2F). Overall, these data demonstrated that in the absence of IL-17A during helminth infection, CD4^+^ T cells in the lung fail to become fully activated and produce type-2 cytokine.

### Neutrophils regulate eosinophil recruitment but not type-2 responses

IL-17A is a major driver for neutrophil recruitment and activation^19^, raising the possibility that impaired neutrophilia in *Il17a-*KO mice is responsible for diminished type-2 immune responses. This hypothesis was supported by evidence that neutrophils from *N. brasiliensis* infected mice can express type-2 cytokines^11^. Therefore, we assessed whether neutrophils contributed to the type-2 immune response by depleting neutrophils during *N. brasiliensis* infection (Fig. 3A). Injection of anti-Ly6G effectively prevented neutrophil accumulation in the BAL at d2pi compared to isotype control (Fig 3B) and depletion was still evident in the blood at d6pi (Suppl. Fig. 2A). Histological sections demonstrated infection-induced injury at d2pi in isotype-treated WT mice, an effect that is almost absent in infected mice depleted of neutrophils. Using lacunarity^20^ as an indicator of acute lung injury^21^, neutrophil depleted mice exhibited no significant signs of damage (Fig. 3C). These data reinforce previous findings that neutrophilia is a major driver of *N. brasiliensis-*induced lung injury^11^. Whilst intestinal worm burdens at d2pi and d4pi were not affected by neutrophil depletion, a small but significant increase in parasite numbers at d6pi was observed in mice treated with neutrophil-depleting antibody (Fig. 3D). As expected, eosinophils were prominent in the lung and BAL of isotype-treated mice at d6pi (Fig. 3E). However, depletion of neutrophils in infected mice significantly reduced eosinophils in the lung and BAL compared to isotype-treated animals. Notably, blood and bone marrow eosinophils were not affected (Supplement Fig. 2B), suggesting a defect in eosinophil recruitment rather than development or survival. We therefore assessed lung mRNA for chemokines with known eosinophil chemotactic effects (*Ccl5, Ccl11*, *Ccl22*), but neutrophil depletion did not alter their expression (data not shown). Because Chen et al. demonstrated that *N. brasiliensis*-primed neutrophils from d2pi exhibit a notable upregulation of *Ccl8*^11^, we assessed expression and found increased *Ccl8* on infection, which was significantly lower in neutrophil-depleted mice (Suppl. Fig. 2C).

**Figure 3:**
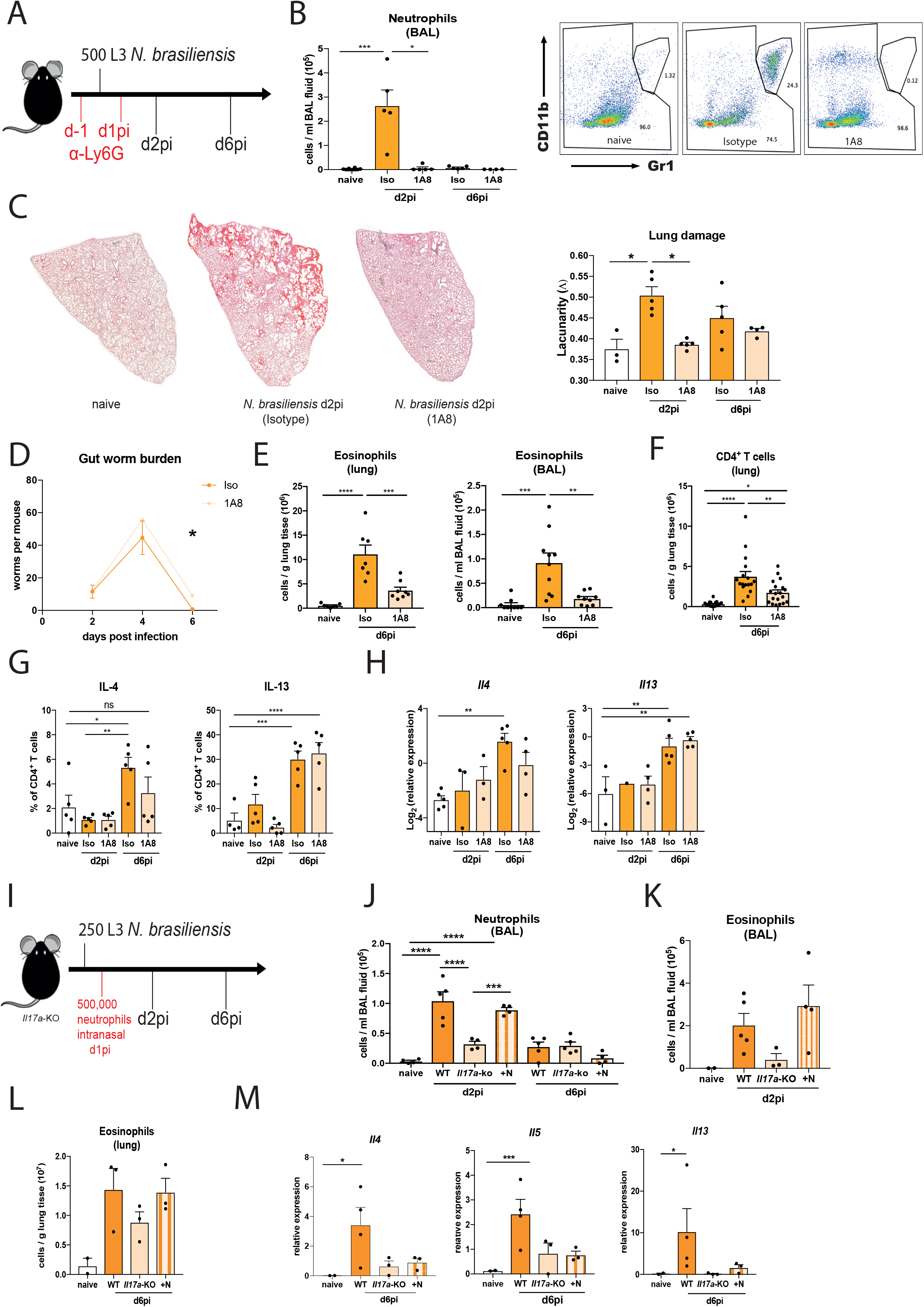
Neutrophils regulate eosinophils but not the type-2 response. C57BL/6J mice were infected with 500 *N. brasiliensis* L3 larvae and mice were injected intraperitoneally with α-Ly6G (1A8) or isotype control on d-1 and d1pi and responses were measured on d2pi or d6pi compared uninfected naïve mice (**A**). Neutrophil numbers per ml BAL, as well as confirmation of neutrophil (Gr1^+^ CD11b^+^) depletion via representative flow cytometry plots (**B**). Representative images of lung sections stained with haematoxylin & eosin and quantification of lacunarity (Λ) on d2pi and d6pi compared to lungs from uninfected naïve mice (**C**). Gut worm burden after neutrophil depletion d2pi, d4pi and d6pi (**D**). Absolute numbers of eosinophils in lung and BAL (**E**), as well as CD4^+^ T cells in the lung (**F**) on d6pi compared to uninfected naïve mice as assessed by flow cytometry. Frequencies of intracellular IL-4^+^ or IL-13^+^ CD4^+^ T cells in lung (**G**). Relative expression of *Il4* and *Il13* in total lung mRNA (log2 expression relative to *actb* (β-actin)) (**H**). *Il17a*-KO mice were intranasally injected with 500,000 neutrophils (+N) d1pi with 250 L3 larvae *N. brasiliensis*, infected controls received PBS, naïve controls were untreated (**I**). BAL neutrophils numbers on days 2 and 6 post infection with *N. brasiliensis* compared to uninfected naïve mice (**J**). Absolute numbers of eosinophils in BAL after neutrophil transfer (**K**). Relative expression of *Il4*, *Il5*, and *Il13* in whole lung mRNA in infected WT *Il17a-*KO or *Il17a-*KO mice + neutrophils (+N) compared to uninfected WT mice (naïve) (**L**). Data are expressed as mean ± s.e.m. and are representative of at least 3 individual experiments with at least 3 mice per group (**B, C, E, G, H, J-L**) or pooled data from two experiments (**D, F**). Data were tested for normality using Shapiro-Wilk test and analysed using one-way ANOVA followed by Sidak’s multiple comparisons test for selected groups. NS – not signification, \**P*<0.05, \*\**P*<0.01, \*\*\**P*<0.001, \*\*\*\**P*<0.0001.

As expected, *N. brasiliensis* infection significantly increased CD4^+^ T cells in the lung, but the number of CD4^+^ T cells were significantly reduced following neutrophil depletion (Fig. 3F). Despite effects on both CD4^+^ T cells and eosinophilia, neutrophil depletion caused limited changes in type-2 cytokine expression (Fig, 3G, H). Although, *Il4* expression or IL-4^+^ CD4^+^ T cells were not significantly upregulated in neutrophil-depleted mice compared to uninfected controls, there were no significant differences detected between neutrophil-depleted versus isotype-treated infected mice (Fig. 3G-H). Additionally, infection-driven increases in IL-13 and IL-5 type-2 cytokines were not altered following neutrophil depletion (Fig. 3G, H, Suppl. Fig. 2D). Thus, neutrophil depletion prevented eosinophil recruitment to the lungs of infected mice but had a limited effect on type-2 cytokine expression.

To further address the role of neutrophils, we tested whether transfer of WT neutrophils into *Il17a-*KO mice could rescue the defect in the development of type-2 immunity. Neutrophils were isolated from the bone marrow of CD45.1^+^ mice and intranasally transferred into *Il17a*-KO mice (Fig. 3I). Neutrophil transfer restored airway neutrophil frequency in *Il17a-*KO mice (Fig. 3J). Interestingly, this neutrophilic response was due to increased recipient-derived cells and not the transferred CD45.1^+^ neutrophils (Suppl. Fig. 2E). The ability of neutrophils themselves to produce neutrophil-active chemokines^22^ may explain this increase in host-derived neutrophils. Regardless of the mechanism, increases to lung neutrophilia one day post-neutrophil transfer (d2pi), rescued eosinophilia in *Il17a*-KO mice, with numbers of eosinophils in IL-17A-deficient animals comparable to infected WT mice (Fig. 3K). *Ccl8* levels were also increased following neutrophil transfer (Suppl. Fig. 2F), consistent with the possibility that CCL8 plays a role during eosinophil recruitment to site of infection. At a later timepoint, when eosinophilia was much more pronounced, the *Il17a-*KO mice that received neutrophils also displayed an increased lung eosinophilia (Fig. 3L). Despite these changes to eosinophil numbers, transfer of neutrophils did not rescue the type-2 cytokine deficit in *Il17a*-KO mice (Fig. 3M). Together, this data provides evidence that neutrophils support eosinophil recruitment at the site of infection but suggests that neutrophils themselves do not directly contribute to pulmonary type-2 cytokine responses.

### IL-17A leads to a downregulation of early IFNγ during *Nippostronglyus* infection

Rapid early IL-17A production is critical for protective immune responses in different settings of lung immunity^1,23^. To better understand the early events unfolding in the lung during *N. brasiliensis* infection, we performed a Nanostring gene expression array using a myeloid immunity panel (700 genes). In whole lung, differentially expressed (DE) genes between naïve WT mice and infected WT and *Il17a-*KO mice at d2pi were assessed in total unamplified RNA (Fig. 4A). IL-17A deficiency led to a distinct gene expression profile compared to WT mice in response to *N. brasiliensis* infection. Notably, when analysing all DE genes (Fig. 4A) using the Ingenuity pathway analyser (Qiagen), IFNγ was predicted as the most significantly increased upstream regulator in *N. brasiliensis* infected *Il17a-*KO compared to WT mice (Fig. 4B). This led us to hypothesize that IL-17A may suppress IFNγ, which would facilitate Th2 cell development and explain why mice deficient in IL-17A cannot induce a full type-2 immune response. This hypothesis was also consistent with our unpublished and published^24^ finding that Ym1, which induces IL-17A, strongly suppresses IFNγ. To test this possibility, we assessed IFNγ responses in *Il17a-*KO mice after *N. brasiliensis* infection. While WT mice exhibited significant suppression of *Ifng* in whole lung at 2dpi compared to uninfected controls, mice deficient in IL-17A did not show this phenotype (Fig. 4C). By intracellularly staining for IFNγ, we observed that *Il17a-*KO mice infected with *N. brasiliensis* failed to exhibit the early downregulation of IFNγ expression seen in infected WT mice (Fig. 4D). Importantly, this failure of suppression was observed across different types of IFNγ-producing cells. Lung CD4^+^ T cells, CD8^+^ T cells, NK1.1^+^ natural killer cells as well as γδ T cells from *Il17a-*KO mice produced IFNγ at the same proportion as the naïve controls, while WT mice exhibited a downregulation in IFNγ production (Fig. 4D). We further assessed whether early IFNγ was produced by γδ T cell subsets that differ in their expression of CD27 and CD44^25^. At 16h post *N. brasiliensis* infection, IFNγ frequencies were also significantly increased in γδ T cells of *Il17a-*KO mice compared to WT controls (Fig. 4E). Consistent with expectations^25^, the CD27^+^ γδ T cells were the main producers of IFNγ after infection (Fig. 4F). Overall, our data demonstrated that IL-17A-deficiency enhanced IFNγ production during infection, supporting the hypothesis that IL-17A plays an important role in downregulating IFNγ at the site of infection during the lung migratory phase of *N. brasiliensis* infection.

**Figure 4:**
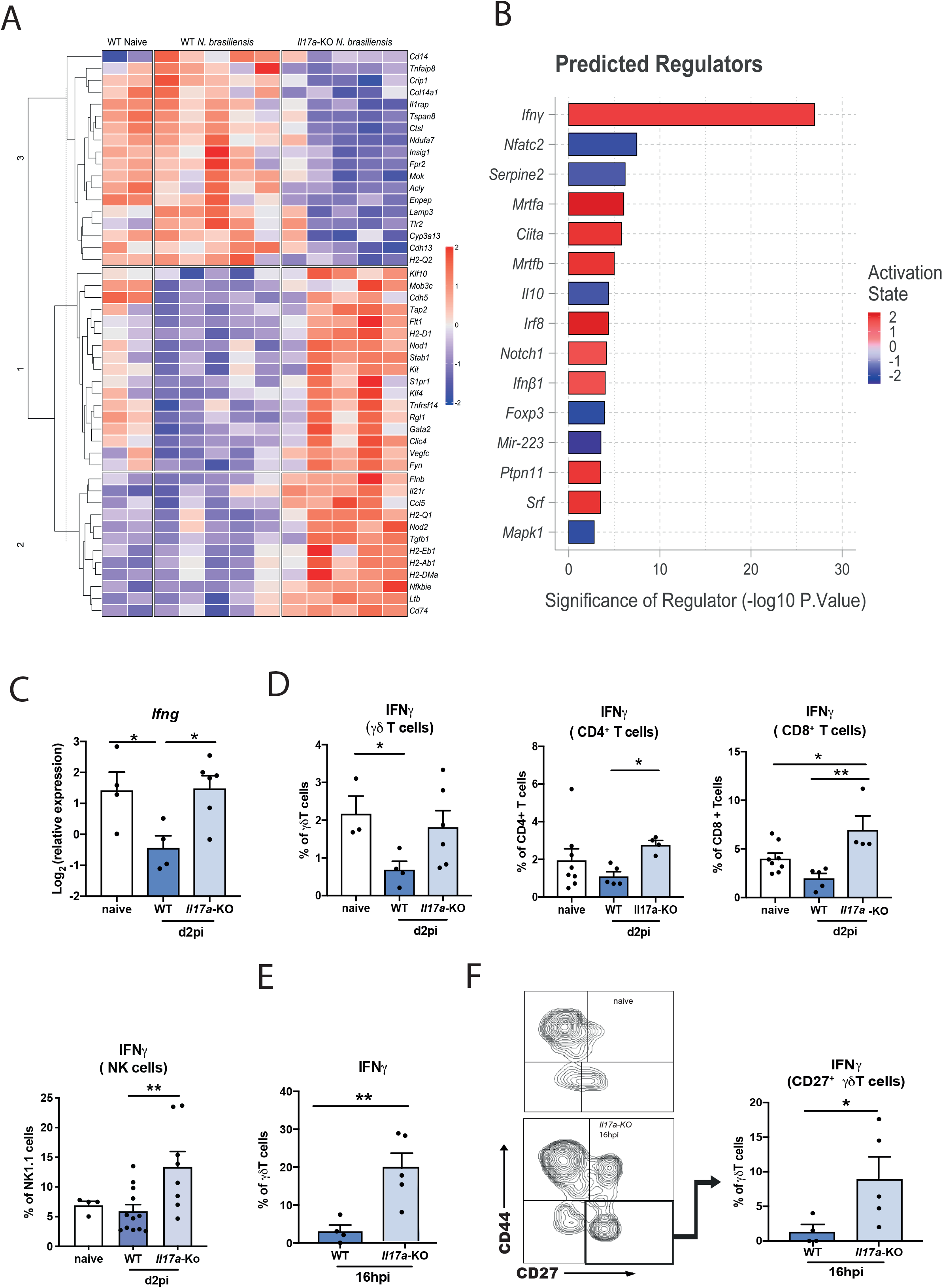
Presence of IL-17A leads to a downregulation of early ifNγ during *N. brasiliensis* infection. Whole lung RNA from C57BL/6J (WT) and *Il17a*-KO mice on d2pi with *N. brasiliensis* compared to WT naïve, were analysed by Nanostring. Unsupervised, hierarchically clustered heat map showing significant differentially expressed genes between infected WT, *Il17a-*KO mice and uninfected (naïve) WT (**A**). Top differentially regulated genes from (**A**) between infected WT and *Il17a-*KO mice were run in Ingenuity pathway analyzer, with top predicted regulators shown in (**B**). Relative expression of *Ifng* in whole lung of naïve WT and d2 *N. brasiliensis* infected WT and *Il17a-*KO mice (log2 expression relative to *actb* (β-actin)) (**C**). Frequencies of IFNγ^+^ γδ T cells, CD4^+^ T cells, CD8^+^ T cells and NK cells in WT and *Il17a-* KO mice d2pi compared to WT naïve mice as assessed by flow cytometry (**D**). Frequency of IFNγ^+^ γδ T cells 16h post *N. brasiliensis* infection in WT and *Il17a-*KO mice (**E**). Representative flow plot showing CD 44 and CD27 γδ T cell subsets in naïve WT mice and *Il17a*-KO mice 16h post *N. brasiliensis* infection as well as frequency of IFNγ^+^ CD27^+^ γδ T cells 16h post *N. brasiliensis* infection in WT and *Il17a-*KO mice (**F**). Data (**C-F**) are expressed as mean ± s.e.m. and are representative of at least 2 individual experiments with at least 3 mice per infected group. Data were tested for normality using Shapiro-Wilk test and analysed using one-way ANOVA followed by Sidak’s multiple comparisons test for selected groups or student’s t-test. \**P*<0.05, \*\**P*<0.01.

### IFNγ neutralization in *Il17a*-KO mice rescues the impaired type-2 immune response

We next asked whether global suppression of early IFNγ by IL-17A was required for the full development of type-2 immunity in the lung. IFNγ was neutralised at day −1 and 1 of infection in *Il17a-*KO and WT mice, and responses examined at d8pi, a time point when the type-2 response should be fully developed (Fig. 5A). The significant defect in eosinophilic responses in *Il17a*-KO mice compared to WT mice was still evident at d8pi. However, blocking IFNγ in *Il17a*-KO mice enhanced eosinophil numbers (Fig. 5B). The same pattern was observed for numbers of CD4^+^ T cells in the lungs (Fig. 5C). To determine whether IFNγ neutralisation altered the type-2 response, we assessed expression of the key type-2 cytokines *Il4* and *Il13* and the type-2 marker, *Chil3*. As expected, based on our results thus far, expression of these genes in the lungs was significantly reduced in infected *Il17a-*KO mice compared to WT mice (Fig. 5D, E). Notably, IFNγ depletion completely recovered expression of these cytokines in *Il17a*-KO mice compared to isotype-treated animals, and *Il4* surpassed the levels seen in WT mice. Similarly, analysis of numbers of IL-5 and IL-13 producing CD4^+^ T cells at d8pi showed restoration of the type-2 response in *Il17a-*KO mice that received the neutralising IFNγ antibody (Fig. 5F). Consistent with the ability of IFNγ to regulate type-2 cytokines, IFNγ depletion also restored the activation status of CD4^+^ T cells in the *Il17a-*KO mice (Fig. 5G) and increased the numbers of CD4^+^ T cells expressing type-2 markers EGFR, PD1 and ST2 (Fig. 5H). Again, no effect was observed in these parameters in IFNγ-depleted WT mice. Together these data demonstrate that an initial reduction in IFNγ levels during *N. brasiliensis* infection mediated by IL-17A, allows the subsequent development of type-2 immunity in the lung.

**Figure 5:**
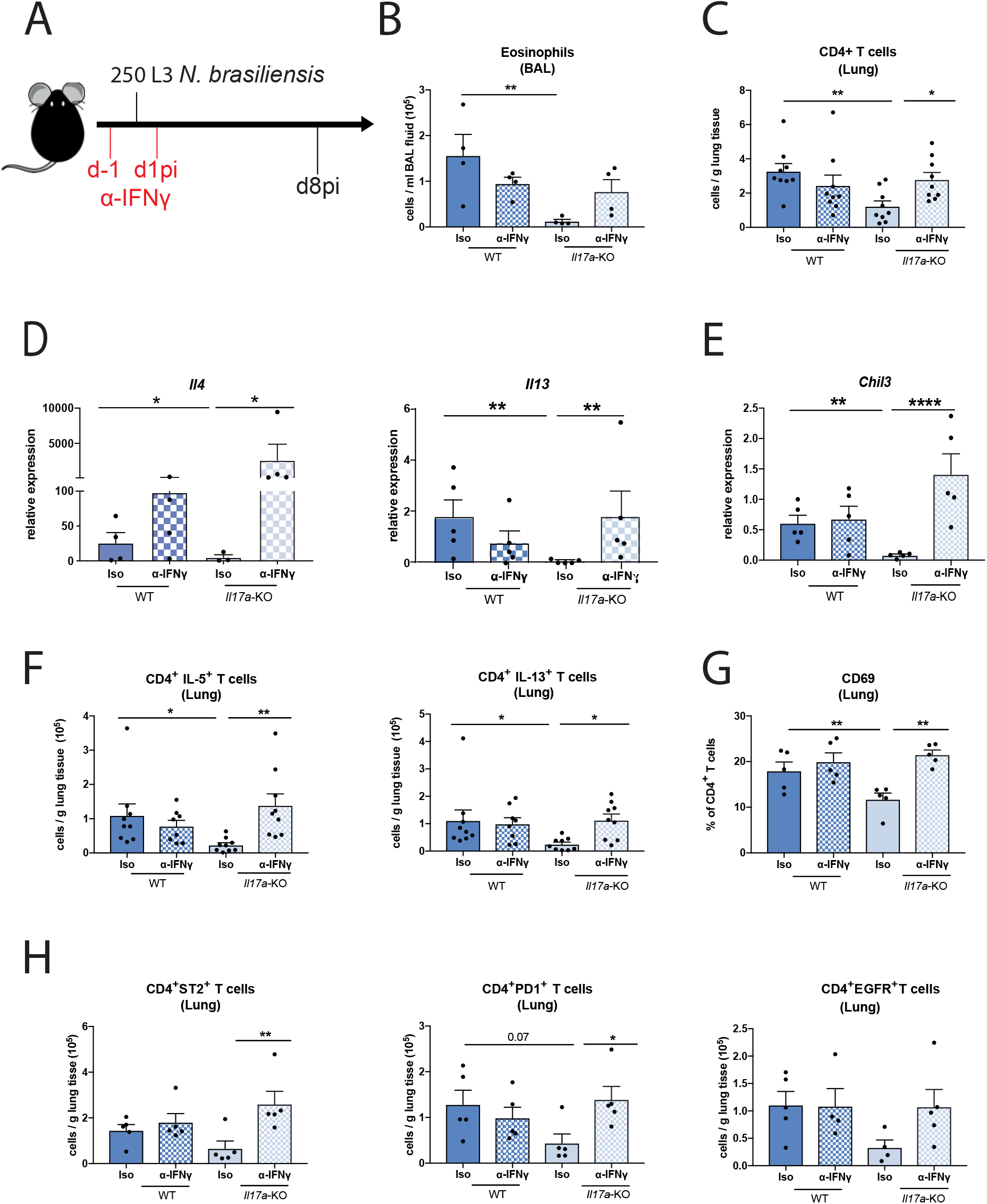
ifNγ neutralization in *Il17a-*KO mice rescues the impaired type-2 immune response. C57BL/6J (WT) and *Il17a-*KO mice and were treated with α-IFNγ or isotype control on days-1 and 1pi with 250 L3 larvae of *N. brasiliensis* (**A**). Absolute numbers of eosinophils per mL of BAL (**B**) or CD4^+^ T cells per gram lung tissue (**C**) as measured via flow cytometry on d8pi. Relative mRNA expression of type-2 cytokines *Il4* and *Il13* (**D**) and type-2 marker *Chil3* (**E**) from whole lung. Absolute numbers of IL-5^+^ and IL-13^+^ CD4^+^ T cells (**F**). Frequency of CD69^+^ CD4^+^ T cells (**G**) and numbers of EGFR^+^, ST2^+^ and PD1^+^ CD4^+^ T cells per gram lung tissue (**H**). Data (**B, D, E, F, G**) are representative (mean ± s.e.m.) of 2 individual experiments with at least 3 mice per group (per experiment) or pooled data from two experiments (**C, F**). Data was tested for normality using Shapiro-Wilk test and analysed using one-way ANOVA followed by Sidak’s multiple comparisons test for selected groups. Data in (**D, E**) were log2 transformed to achieve normal distribution and statistical tests were performed on transformed data. \**P*<0.05, \*\**P*<0.01, \*\*\**P*<0.001.

### IL-17A suppresses an established type-2 response in the lung

IFNγ depletion in *Il17a-*KO mice not only restored the type-2 response, but in some cases exceeded WT levels. Therefore, we hypothesized that although innate IL-17A promotes the establishment of type-2 immunity in *N. brasiliensis* infection, once the adaptive response is in place, IL-17 can act to negatively regulate the type-2 pulmonary response. To test this hypothesis, we neutralized IL-17A at d4pi, d5pi and d6pi in WT mice and assessed immune responses at d7pi (Fig. 6A). Blocking of IL-17A led to a significant increase in both ILC2 numbers and frequencies in the lung (Fig. 6B), as well as the numbers of ILC2s producing IL-5 and IL-13 (Fig. 6C). Although CD4^+^ T cell numbers in the lung were comparable between isotype-treated and anti-IL-17A-treated WT mice (Fig. 6D), the ability of CD4^+^ T cells to produce type-2 cytokines may partly rely on IL-17A, as mice administered anti-IL-17A showed a slight increase IL-5 and IL-13 (Fig. 6E). This data demonstrated that IL-17A can have differential effects depending on the time and status of infection. While early IL-17A promotes the type-2 response, later in infection IL-17A acts to suppress and limit excessive the type-2 immunity, particularly in ILC2s.

**Figure 6:**
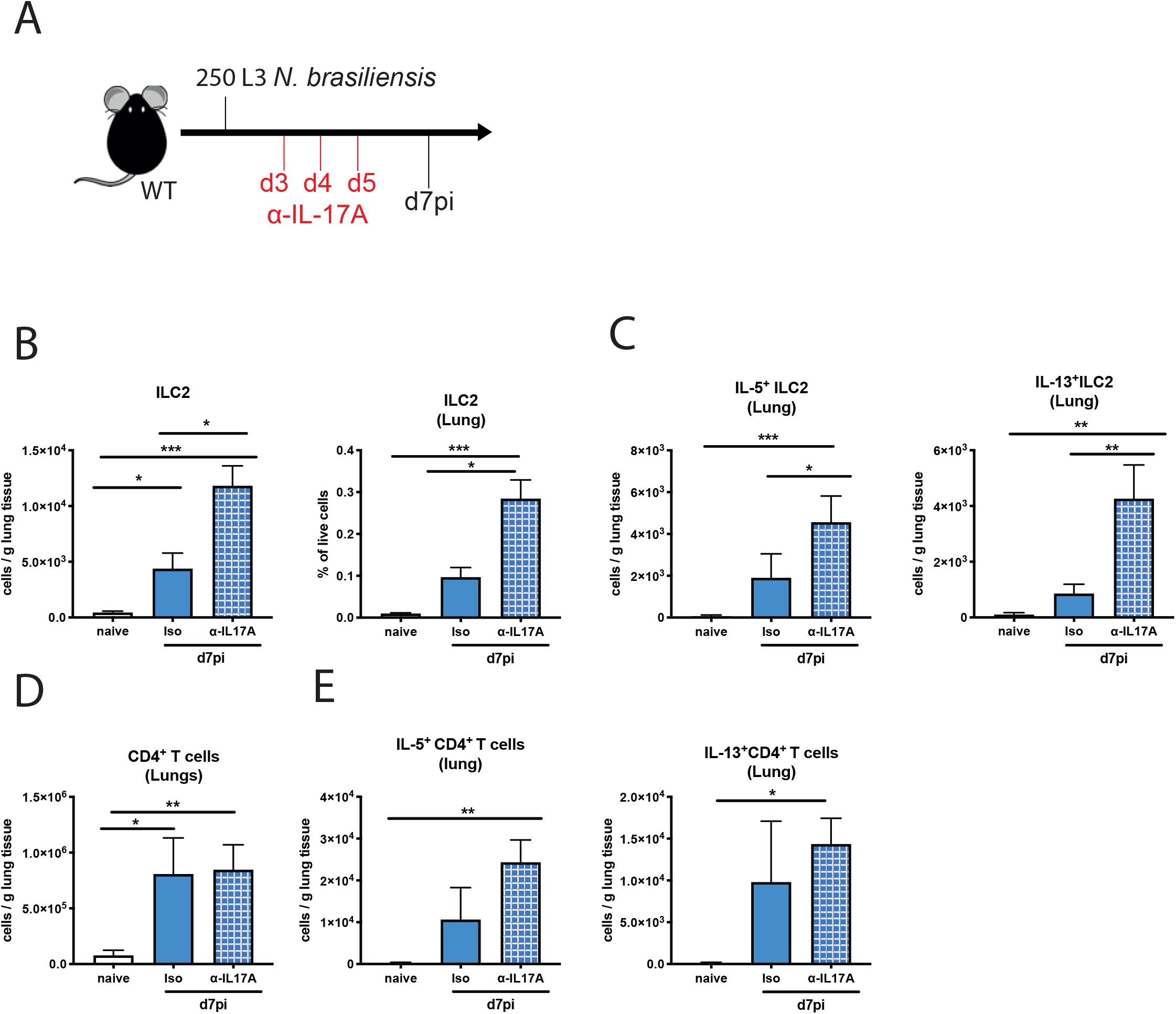
Late stage IL-17A suppresses type-2 immune responses during *N.brasiliensis* infection. C57BL/6J WT mice were treated with α-IL-17A or isotype control on days 3, 4, and 5pi with 250 L3 *N. brasiliensis* and responses measured at d7pi compared to uninfected (naïve) mice (**A**). Cells per gram lung tissue and frequency of ILC2 in live lung cells (**B**). Absolute number of IL-5^+^ and IL-13^+^ ILC2 per gram lung tissue (n=5 for naïve, n=11-12 for d7 *N. brasiliensis* infected groups) (**C**). Absolute numbers of CD4^+^ T cells (**D**) and IL-5^+^ and IL-13^+^ CD4^+^ T cells per gram of lung tissue (n=3 for naïve and n=6 for d7 *N. brasiliensis* infected groups) (**E**). Data pooled from two independent experiments (**B**, **C**) or are representative (mean ± s.e.m.) of 3 individual experiments with at least 3 mice per group (per experiment) (**D**, **E**). Data was tested for normality using Shapiro-Wilk test and analysed by a one-Way ANOVA followed by Sidak’s multiple comparisons test for selected groups. \**P*<0.05, \*\**P*<0.01 \*\*\**P*<0.001.

### IL-17A does not regulate type-2 immune responses at the site of *T. muris* infection

Our data demonstrate an impairment of the type-2 immune response in the lung during infection *Il17a-*KO mice with the lung-migrating nematode *N. brasiliensis*. We wanted to investigate whether impairment of type-2 immunity by IL-17A was unique to the pulmonary environment. Initially we examined type-2 cytokine gene expression in the small intestine of *N. brasiliensis* infected mice at d7pi. However, we did not observe any significant changes in WT infected mice (data not shown). We therefore decided to use *Trichuris muris,* a nematode that establishes infection solely in the gastro-intestinal tract. Infection with *T. muris* begins with the ingestion of infective eggs that accumulate in the caecum. L1 larvae hatch and penetrate the caecum and proximal colon wall, undergoing moults to L2 (d9-11pi), L3 (d17pi), L4 (d22 pi) and adults (d29-32). High dose infection of C57BL/6 mice induces a strong type-2 response by d17pi, and subsequent clearance of the adult parasites^26,27^. We infected WT and *Il17a-*KO mice with a high dose of 200 *T. muris* eggs and found worm counts in the caecum were comparable at d19pi and d32pi (Fig. 7A), indicating IL-17A does not alter parasite expulsion rate. Cell numbers in the caecum were analysed and no differences in eosinophil and neutrophil frequency were observed between *Il17a-*KO mice and WT controls on d19pi and d32pi (Supplement Fig. 3A, B), suggesting an IL-17A-independent recruitment mechanism for both these cell types. CD4^+^ T cell numbers in the mLN were also comparable on d19pi and d32pi between *Il17a-*KO mice and WT controls (Suppl. Fig 3C). Although there was an induction of type-2 cytokines in infected mice as measured by intracellular cytokine staining, the numbers of IL-4, IL-5 or IL-13-producing CD4^+^ T cells in the MLNs did not significantly differ between the groups (Suppl. Fig 3D). Similarly, the relative expression of the cytokines *Il4*, *Il5* and *Il13* did not differ between *Il17a-*KO mice and WT controls within the caecum (Suppl. Fig 3E). Secreted levels of IL-5, IL-9 and IL-13 in MLN cells were also not impaired in *Il17a-*KO mice relative to WT controls (Suppl. Fig 3F). Together, these data failed to provide any evidence that IL-17A was an important regulator of type-2 immunity in the intestine during *T. muris* infection.

**Figure 7:**
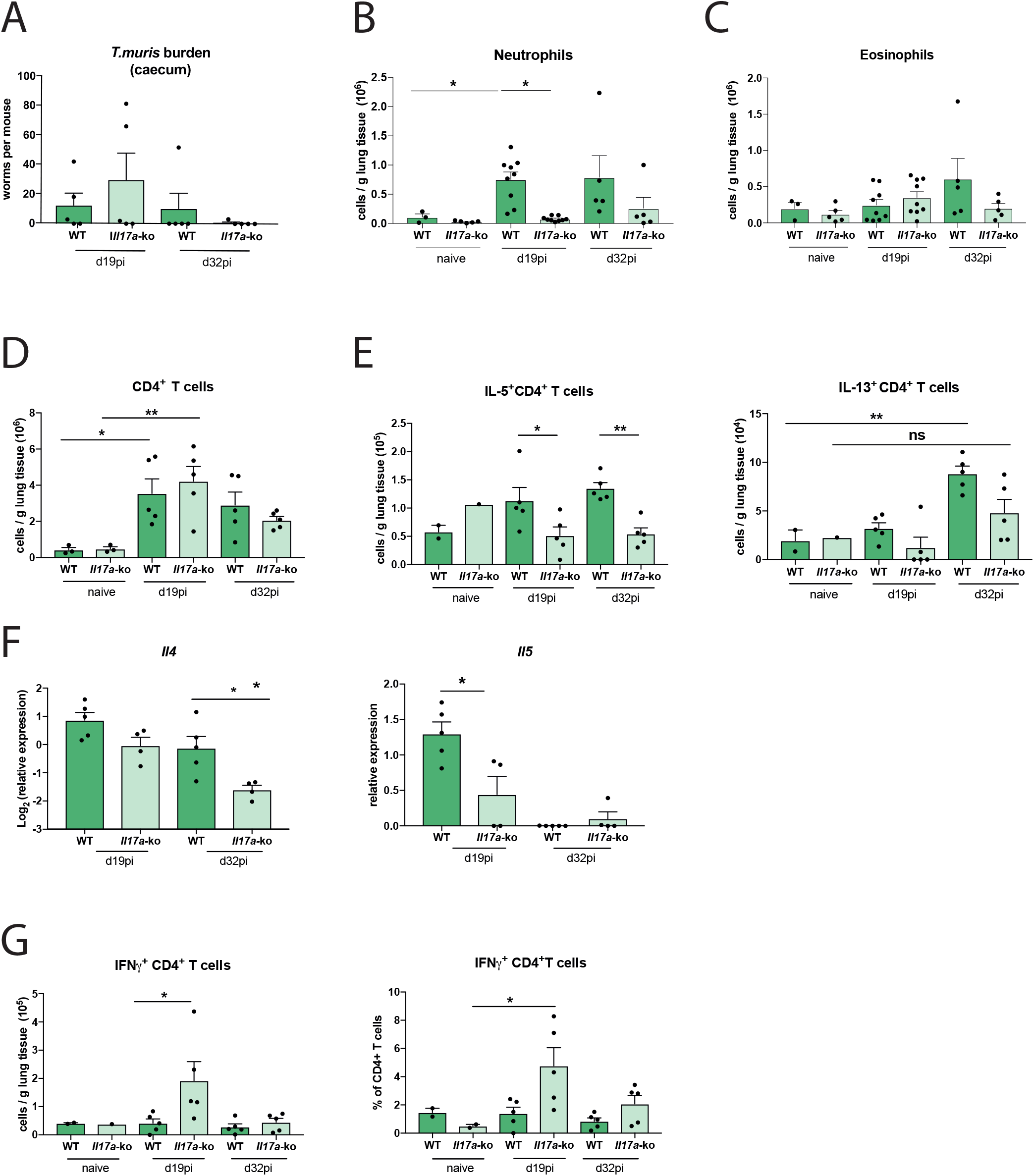
Lack of IL-17A impairs concurrent type-2 immune responses in the lung following infection with *Trichuris muris*. C57BL/6 WT and *Il17a*-KO mice were infected with a high dose of *T. muris* and immune parameters investigated at d19 and d32pi compared to uninfected (naïve) C57BL/6 WT and *Il17a*-KO mice. Worms counts in the caecum (**A**). Absolute numbers of neutrophils (**B**), eosinophils (**C**) and CD4^+^ T cells (**D**) per gram of lung at d19pi and d32pi compared to naïve mice. Absolute numbers of IL-5^+^ and IL-13^+^ CD4+ T cells per gram of lung tissue (**E**). Relative mRNA expression of cytokines *Il4* and *Il5* from whole lung (log2 expression relative to *actb* (β-actin)) of infected mice (**F**). Absolute numbers and frequency of IFNγ^+^ CD4^+^ T cells per gram of lung tissue (**G**). Data are expressed as mean ± s.e.m. and are representative of 3 individual experiments with at least 4 mice per infected group and one mouse per control group. Data was tested for normality using Shapiro-Wilk test and analysed with one-way ANOVA followed by Sidak’s multiple comparisons test for selected groups. \**P*<0.05, \*\**P*<0.01

Previous studies have shown that despite the restriction of the *T. muris* lifecycle to the gastro-intestinal tract of the mammalian host, evidence of a type-2 immune response can be observed at distant sites, such as the lung^28^. Therefore, the immune response in the lung of *T. muris* infected WT vs *Il17a-*KO mice at d19pi and d32pi was assessed. Neutrophils numbers were increased in infected WT animals at d19pi and d32pi, but this was significantly reduced in *Il17a-*KO mice on d19pi (Fig. 7B). No significant changes were observed for eosinophils (Fig. 7C). Whilst lung CD4^+^ T cell numbers in infected animals did not change compared to naïve controls (Fig. 7D), *Il17a*-KO mice had significantly fewer IL-5^+^CD4^+^ T cells at d19 and d32pi compared to WT controls (Fig. 7E). Although the effect on IL-13^+^CD4^+^ T cells was less evident, infected *Il17a*-KO mice failed to significantly increase numbers of IL-13^+^CD4^+^ T cells compared to uninfected controls (Fig. 7E). Supporting the intracellular cytokine staining, qRT-PCR analysis in whole lung tissue showed an impairment of type-2 cytokines in the *Il17a-*KO mice, with significantly decreased expression of *Il4* (d32pi) and *Il5* (d19pi) (Fig. 7F).

Similar to infection with *N. brasiliensis*, we also observed an upregulation of IFNγ in the lung during *T. muris* infection in *Il17a-*KO mice. Both the number and the frequency of IFNγ^+^CD4^+^ T cells in the lung were significantly increased in *Il17a-*KO compared to WT infected mice on d19pi (Fig. 7G). This data utilising *T. muris* infection models suggests that IL-17A-dependent suppression of IFNγ allows promotion of the type-2 immune response specifically in the lungs but not the intestine and highlights a means of communication between the intestine and the lung involving IL-17A, not previously described.

## DISCUSSION

IL-17A, the key cytokine of the IL-17 family, is central to barrier immunity, combating fungal infections and inducing antimicrobial proteins as well as neutrophil activating and recruiting chemokines^2^. However, in the context of type-2 immunity, a combination of type-2 cytokines and IL-17A is often a signature for severe disease pathology. For example, IL-17A contributes to asthma pathology by enhancing IL-13 activity^29^ and a dysregulated balance between IL-17A and type-2 responses exacerbates pathology during schistosomiasis and onchocerciasis^4–6,30^. Understanding the relationship between IL-17A and type-2 immune responses is thus critical, and we and others have previously demonstrated that development of a full type-2 response can require IL-17A^8,12,13,31^.

In our effort to understand how IL-17A might be required for full type-2 immunity, we have discovered that IL-17A suppresses early IFNγ expression in the lung during helminth infection. Although several studies show links between IL-17A and IFNγ, whether IFNγ is up- or downregulated in response to IL-17A varies with setting, timing and location. For example enhanced IFNγ in *Il17a-*KO mice has been described in a viral infection^32^, experimental visceral leishmaniasis, and *Toxoplasma gondii* infection^33^. Evidence also exists in the context of helminth infection, where a lack of IL-17A drives elevated IFNγ during infection with the filarial nematode *Litomosoides sigmondontis*^34^ or *Schistosoma japonicum*^35^ and *Schistosoma mansoni*^5^. In contrast, IL-17A can promote IFNγ production during kidney-ischemic reperfusion injury^36^, or *Francisella tularensis* infection^37^. Importantly, the consequence of IL-17A-IFNγ cross-regulations in the context of type-2 inflammation has never been shown and here we reveal IFNγ downregulation as a new mechanism through which IL-17A establishes a protective type-2 response in the lung.

Another key finding of our study was that the requirement for IL-17A to suppress IFNγ appears restricted to the lung. The protective type-2 immune response in the gut of *Il17a-*KO mice was not impaired, and mice were still able to expel *N. brasiliensis* from the small intestine and *T. muris* from the colon. The more surprising finding was that even though *T. muris* does not have a lung stage, the concurrent type-2 response in the lung was impaired in *Il17a-*KO mice. CD4^+^ T cells in the lung produced less type-2 cytokines and both eosinophils and neutrophils were significantly downregulated in the *Il17a-*KO mice. Consistent with our findings in *N. brasiliensis*, CD4^+^ T cells in *Il17a-KO* mice produced significantly higher amounts of IFNγ than in their WT counterparts, raising major questions as to the nature of the insult that induces IL-17 in the lungs of *T. muris* infected mice. Although our data suggest that the impact of IL-17A on type-2 development may be lung restricted, there may still be a fundamental requirement for suppression of IFNγ for type-2 immunity to progress. Artis et al. demonstrated that the type-2 immune response during *T. muris* requires TSLP, and in very similar experiments to those described here, demonstrated that TSLP functions to suppress IFNγ^38^. Thus, early suppression of IFNγ may be a general pre-requisite for the development of a type-2 environment with a requirement for IL-17A in the lung and TSLP (or other factors) in the gut.

We have not yet addressed the full mechanism behind IL-17A-mediated suppression of IFNγ during *N. brasiliensis* infection but it is notable that IL-17A not only impairs type-2 cytokine production, but also alters the cellular activation status and expression of type-2 markers. Interestingly, in our model, we only observe impairment of type-2 immune responses in the lung itself and not in the Th2 cells from lung-draining lymph nodes. Expression of EGFR and ST2, two markers closely associated with type-2 settings^16^, were reduced on the CD4^+^ T cells of *Il17a-*KO mice in the lung. EGFR expression on Th2 cells is critical for resistance during GI helminth infection and a signalling complex between EGFR and ST2 can activate Th2 cells to secrete IL-13 in an antigen-dependent manner upon IL-33 exposure. Our data would suggest that this “licensing” of Th2 cells does not occur in the *Il17a-*KO mice during *N. brasiliensis* infection, indicating that IL-17A is needed for a proper induction of the adaptive Th2 response in the lung.

As expected, IL-17A was important for innate neutrophilic responses during *N. brasiliensis* infection. However, neutrophil depletion and transfer experiments demonstrated that neutrophils themselves only partly contributed to the development of type-2 immunity, mainly through enhancement of the subsequent eosinophilia. Our data highlight a potential role for the chemokine CCL8 in neutrophil mediated recruitment of eosinophils. Although CCL8 has been previously associated with eosinophils^39^ and type-2 inflammatory responses^40^ in the lung, neither study demonstrated a direct eosinophil chemotactic function of CCL8. More recently, Puttur et al. revealed that CCL8 is important for the appropriate mobilisation of pulmonary ILC2s and their localised production of IL-5, possibly explaining why we observed reduced eosinophilia in neutrophil depleted mice^41^.

It is well documented that type-2 responses are essential to limit excessive IL-17^42–44^ but a novel finding from our study is that the reverse is also true. While the early γδ T cell-derived IL-17A supported the type-2 response, late IL-17A, derived from both Th17 cells and γδ T cells, negatively regulated type-2 cytokines. To our knowledge, IL-17A suppression of type-2 cytokines has not previously been described in vivo and illustrates a major cross-regulatory axis between type-2 cytokines and IL-17A, each required to contain the excessive production of the other. In the numerous situations in which a combination of type-2 cytokines and IL-17A results in severe disease pathology^4,6,29^, it is apparent that this cytokine balance has failed. Together our data demonstrate that early events in the lung shape the protective type-2 immune response, with IL-17A as a critical regulator of type-2 immunity. IL-17A, as a driver of tissue damage^8^, may itself be needed to establish a subsequent type-2 repair response. However, the ability of IL-17A to then suppress type-2 responses, reveal an important feedback loop that must go awry during severe asthma and other type-2 conditions in which IL-17A plays a damaging and pathogenic role. Finally, in combination with previous data^38^, suppression of IFNγ at barrier sites may be a central paradigm for type-2 immunity.

## MATERIALS AND METHODS

### Mice and ethics statement

For experiments using only WT mice, C57BL/6 J mice were obtained from Charles River. C57BL/6 *Il17a*^Cre^*Rosa26*^eYFP^ mice were originally provided by Dr Brigitta Stockinger^45,46^. For *Il17a-*KO experiments C57BL/6 WT mice and C57BL/6 *Il17a*^Cre^*Rosa26*^eYFP^ homozygote mice were bred at the University of Manchester. Mice were age- and sex-matched and all mice were housed in individually ventilated cages. Both males and females were used. Mice were not randomized in cages, but each cage was randomly assigned to a treatment group. Mice were culled by asphyxiation in a rising concentration of CO_2_. Experiments were performed in accordance with the United Kingdom Animals (Scientific Procedures) Act of 1986.

#### *N. brasiliensis* infection

*N. brasiliensis* was maintained by serial passage through Sprague-Dawley rats, as described^47^. Third-stage larvae (L3) were washed ten times with PBS (Dulbecco’s PBS, Sigma) before infection. On day 0, mice were infected subcutaneously with 250 or 500 larvae (L3). At various time points mice were euthanised, bronchoalveolar lavage (BAL) was performed with PBS containing 1% BSA and lungs were taken for further analysis. For worm counts, the small intestines of infected mice were collected in PBS. Small intestines were then cut longitudinally along the entire length, placed in a 50 ml Falcon and incubated at 37°C for 4h. Settled worms were then counted with the aid of a dissecting microscope.

### Flow cytometry

Single-cell suspensions of the lung were prepared by digesting minced lung lobes for 30 min at 37°C with 0.2 U/ml Liberase TL (Roche) and 80 U/ml DNase (Life Tech) in Hank’s balanced-salt solution before forcing tissue suspensions through a cell strainer (70 μm, Greiner). Red blood cells were lysed using Red Blood Cell Lysing Buffer Hybri Max (Sigma) for 3 min at RT and reaction was stopped by diluting samples in PBS. Total live cells were counted with AO/PI dye on an automated cell counter (Auto2000, Nexcelom). Cells were stained for live/dead (Life Technologies) and then incubated with Fc-block (1:500 CD16/CD32 and 1:50 mouse serum) and were then stained with fluorescence-conjugated antibodies. Cells were identified by expression of surface markers as follows: neutrophils Ly6G^+^CD11b^+^, eosinophils CD11b^+^ CD11c^−^ SigF^+^, CD4 T cells CD4^+^, TCRβ^+^CD11b^−^, γδ Tcells TCRβ^−^, TCRγδ^+^, CD11b^−^ and ILC2s Lineage^−^ (CD11b, TCRβ, TCRγδ, Ly6G, F4/80, CD11c, SigF, CD19) CD90.2^+^KLRG^+^CD127^+^. Antibody clones used are listed in **Table 1**. For staining of intracellular cytokines, cells were stimulated for 4 h at 37°C with cell stimulation cocktail containing protein transport inhibitor (eBioscience), then stained with live/dead. After surface antibody staining, cells were fixed for 10 min at 4°C using IC fix (Biolegend) and cells were then incubated in for 20 min at RT in Permeabilization buffer (biolegend). Intracellular staining was performed for cytokines using antibodies for IL-5, IL-13, IL-17A and IFNγ as well as for Gata3, Ym1 and Relm-α. Samples were analysed by flow cytometry with LSR Fortessa or LSR II (Becton-Dickinson) and data analysed using FlowJo v10 software.

**Table 1.**
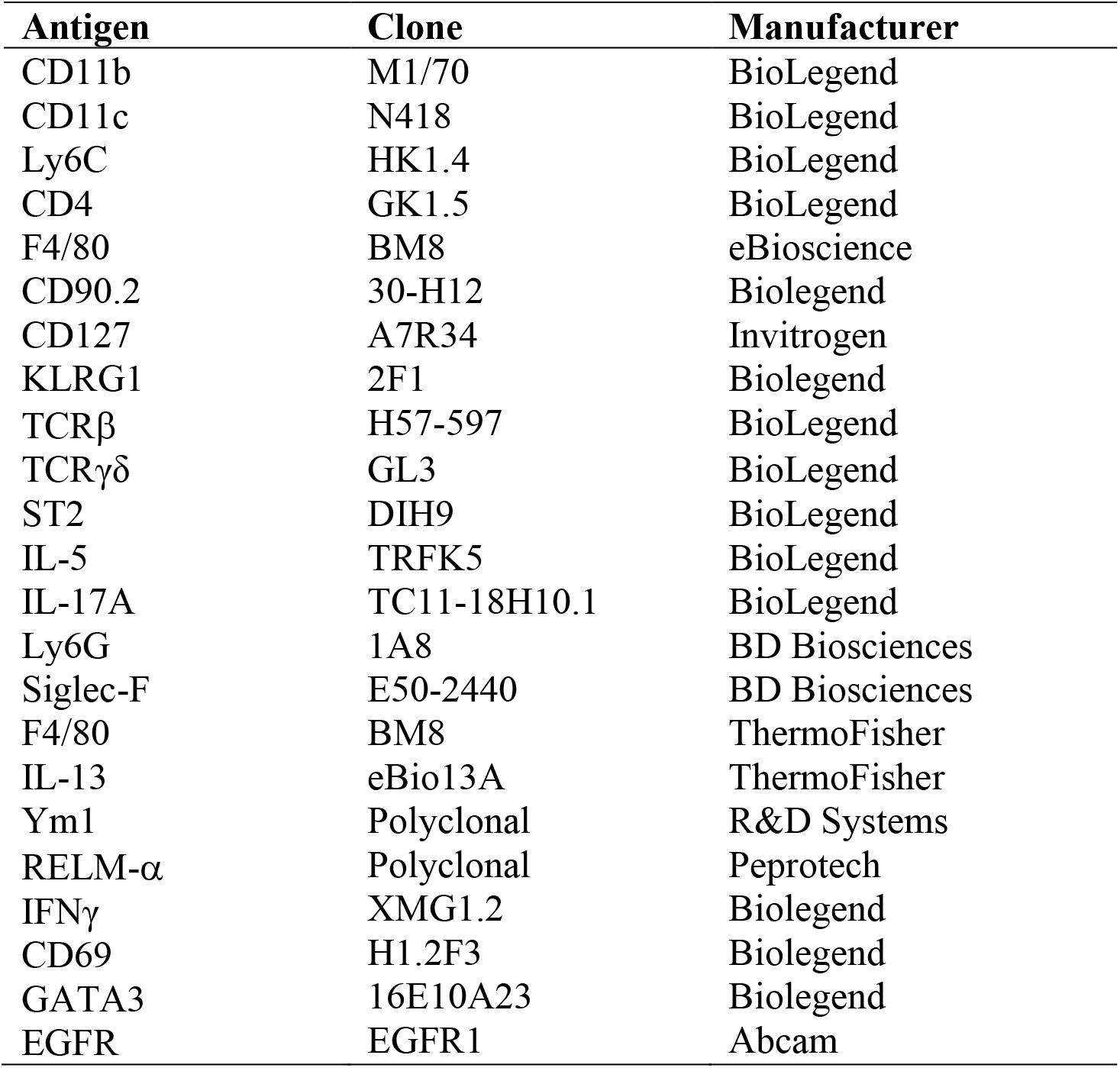
List of flow cytometry antibodies used.

### Quantification of cytokines

Single-cell suspensions of splenocytes, lung-draining lymph nodes or whole lung were stimulated *ex vivo* with *N. brasiliensis* excretory secretory product (E/S) antigen^48^ (1 μg/ ml) or anti-CD3 (1 μg/ml). Cell supernatants were harvested 72 h later and were stored at −20°C until further analysis. Mouse IL-13 DuoSet ELISA kit (R&D Systems) was used for measurement of IL-13 levels. Mesenteric lymph node (MLN) cells from *T. muris* infected or uninfected mice were collected, cultured and restimulated *ex vivo* for 36 h with E/S as previously described ^26^. The concentrations of IL-5, IL-6, IL-9, IL-10, IL-13, IL-17A, TNFα and IFNγ in the mLN culture supernatant were measured by cytokine bead array (CBA, BD Biosciences, UK) as per the manufacturer’s protocol.

### Antibody depletion and cell transfer experiments

Neutrophils were depleted via intraperitoneal injection of anti-Ly6G antibody (clone 1A8, BioXcell, West Lebanon, NH, USA) (500 μg/mouse/day) on days −1 and 1 of infection with *N. brasiliensis*. Control mice were injected with an equal volume of IgG2a isotype control (BioXcell). Neutrophil depletion was confirmed by anti-Gr1 (RB6-8C5) staining for flow cytometry. For neutrophil transfer, neutrophils were purified from bone marrow using Histopaque-based density gradient centrifugation as described before^49^. Collected neutrophils were washed twice with RPMI 1640 and counts and viability was determined (average purity 85% ± 4.42% and with average viability of 90% ± 1.5%). Neutrophils were resuspended in PBS and 0.5 × 10^6^ cells (40 μl) transferred intranasally into mice on day 1 post infection with *N. brasiliensis*. IFNγ was depleted using an anti-IFNγ monoclonal antibody (clone XMG1.2) and injected intraperitoneally (500μg/mouse/day) on days −1 and 1 of infection with *N. brasiliensis*. Control mice were injected with an equal amount of corresponding isotype control (GL113). IL-17A was depleted using an anti-IL-17A (17F3) or IgG1 isotype (both Invivo mAB) injected intraperitoneally (100μg/mouse/day) on days 4, 5 and 6 post-infection with *N. brasiliensis*.

### Histology

For histology, the left lobe of the lung was isolated at different time points following *N. brasiliensis* infection, inflated and fixed in 10 % neutral-buffered formalin (Sigma). Whole left lung lobes were processed using a tissue processor (Leica ASP300S) and were then embedded in paraffin. Paraffin blocks were then sectioned to 5 μm using a microtome (Leica RM2235) and routinely stained with Mayer’s haematoxylin (Merck Millipore HX86014349) and eosin (Sigma) for histological analysis. Slides were imaged using an Olympus slide scanner and high-resolution image files were exported using Pannoramic Viewer software (3DHISTECH). The images were then processed in a KNIME software workflow to obtain 50 random regions of interests (ROIs) across the whole lung section. ROIs that contained lobe boundaries or extensive artefacts were excluded from the analysis. The ROIs were then converted to binary images and lacunarity (Λ) was quantified using the FracLac plugin for ImageJ^21^. The Λ values of all the ROIs were averaged to obtain estimates for the entire lobe.

### Extraction of RNA and quantitative real-time PCR

A fragment of the right lung lobe was stored in RNAlater (Ambion) before homogenization of tissue in Qiazol reagent with a TissueLyser (Qiagen). RNA was prepared according to manufacturer’s instructions. RNA was quantified using a ND-1000 Spectrophotometer (NanoDrop Technologies). Reverse transcription of 1 μg of total RNA was performed using Tetro reverse transcriptase (Bioline). For reverse transcription, total RNA was treated with 50 U Tetro reverse transcriptase (Bioline), 40 mM dNTPs (Promega), 0.5 μg PolyT primer for cDNA synthesis (Roche) and RNasin inhibitor (Promega). The abundance of transcripts from the genes of interest was measured by quantitative real-time PCR with the Light Cycler 480 II system (Roche) with a Brilliant III SYBR master mix (Agilent) and specific primer pairs. PCR amplification was analysed by the second-derivative maximum algorithm (Light Cycler 480 Sw 1.5; Roche), and expression of the gene of interest was normalized to that of the housekeeping gene *Actb* (beta-actin). A list of primer sequences used are shown in **Table 2**.

**Table 2.**
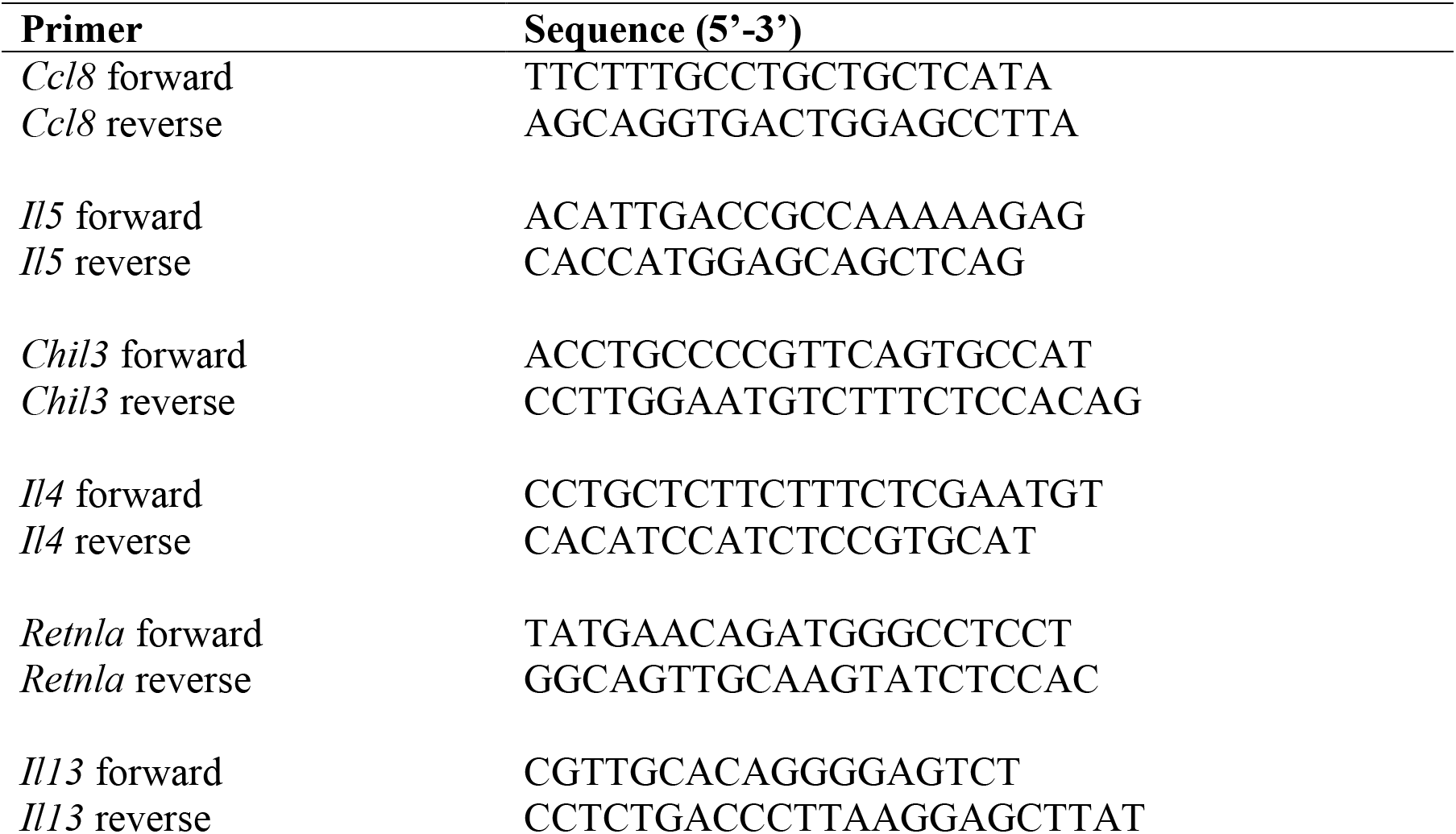
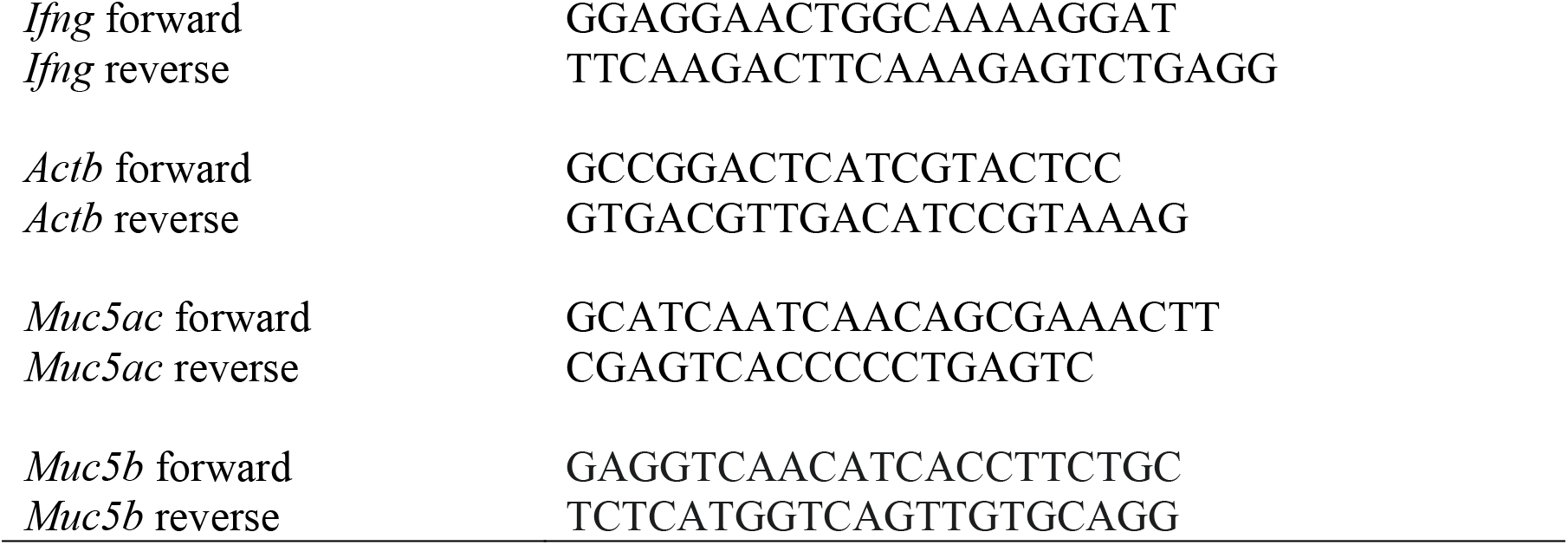
List of primer sequences used.

### *Trichuris muris* infection and E/S products

*T. muris* eggs were prepared from chronically infected stock mice as described previously^50^. Mice were infected by oral gavage with 200 embryonated *T. muris* eggs suspended in ddH2O. At day 19 and 32 post infection, *T. muris* burden was assessed by removing the caecum and proximal colon, opening them longitudinally and scraping the contents out with fine forceps. Individual worms were then counted by eye under a binocular dissecting microscope. *T. muris* adult excretory secretory product antigen (E/S) was prepared as described by^50^. In brief, adult *T. muris* were cultured *ex vivo* at 37°C, the culture supernatant was collected and centrifuged to remove eggs and worms. The resultant supernatant was then filter sterilised and stored at −20°C until use for *in vitro* re-stimulation of MLN cells.

### Nanostring RNA Profiling

Extracted RNA was run on an Agilent 2200 Tape Station system to ensure high quality lung RNA; samples with a RIN value of < 6.5 were excluded. Suitable RNA was then diluted to 20 ng/μL in RNase free H2O, measured using Qubit™ RNA HS Assay Kit (Thermofisher) and run on a Nanostring nCounter® FLEX system using the Myeloid Innate Immunity v2 panel (XT-CSO-MMII2-12) 220 as per manufacturer’s instructions. Raw data was loaded into nSolver version 4.0 using default settings. Non-normalised counts were then exported and subsequent analyses were performed in R (version 3.6) using RStudio Version 1.2.1335 Build 1379 – © 2009-2019 RStudio, Inc. Positive controls were analysed to ensure there was clear resolution at variable expression levels and negative controls were used to set a minimum detection threshold which was applied to all samples. Data were then normalised with EdgeR using the TMM method and differential expression between *N. brasiliensis*-infected WT and *Il17a*-KO mice was calculated via linear modelling with Empirical Bayes smoothing using the limma R package 2^51^. Genes with an absolute fold change of greater than one and a significance value of under 0.05 after correction for multiple comparisons using the Benjamini-Yekeuteli method were defined as ‘differentially expressed’ and taken forward for further analysis. Heatmaps were then generated from normalized counts of differentially expressed genes using the ComplexHeatmaps R package. The networks and functional analyses of differentially expressed genes were generated with Ingenuity pathway analyser (QIAGEN Inc., https://www.qiagenbio-informatics.com/products/ingenuity-pathway-analysis). Data were then imported into R for visualisation.

### Statistics

Prism 7.0 (version 7.0c, GraphPad Software) was used for statistical analysis. Differences between experimental groups were assessed by ANOVA (for normally distributed data, tested using Shapiro-Wilk normality test) followed by Sidak’s multiple comparisons test. For gene expression data, values were log_2_ transformed to achieve normal distribution. Comparisons with a *P* value of less than 0.05 were considered to be statistically significant. Data is represented as mean ± sem.

## ACKNOWLEDMENTS

We thank the Flow Cytometry, Bioimaging, and Biological Services core facilities at the University of Manchester. This work was supported by the Wellcome Trust (106898/A/15/Z to JEA and Z10661/Z/18/Z to RG), the Medical Research Council UK (MR/K01207X/1 to JEA), Medical Research Foundation UK joint funding with Asthma UK (MRFAUK-2015-302 to TES). SC was supported by a Wellcome Trust Studentship (103132/Z/13/Z). We thank Kevin Couper for CD45.1 mice.

## AUTHOR CONTRIBUTION

J.A., A.L.C., J.E.P and S.A.P.C. executed experiments, B.H.K.C. and S. P. provided experimental assistance. J.A., R.K.G., T.E.S. and J.E.A. designed experiments and analysed data. L.B. provided reagents. J.A., T.E.S. and J.E.A. wrote the original draft of the manuscript and all co-authors reviewed and edited the manuscript.

---

**Supplement Figure 1:**
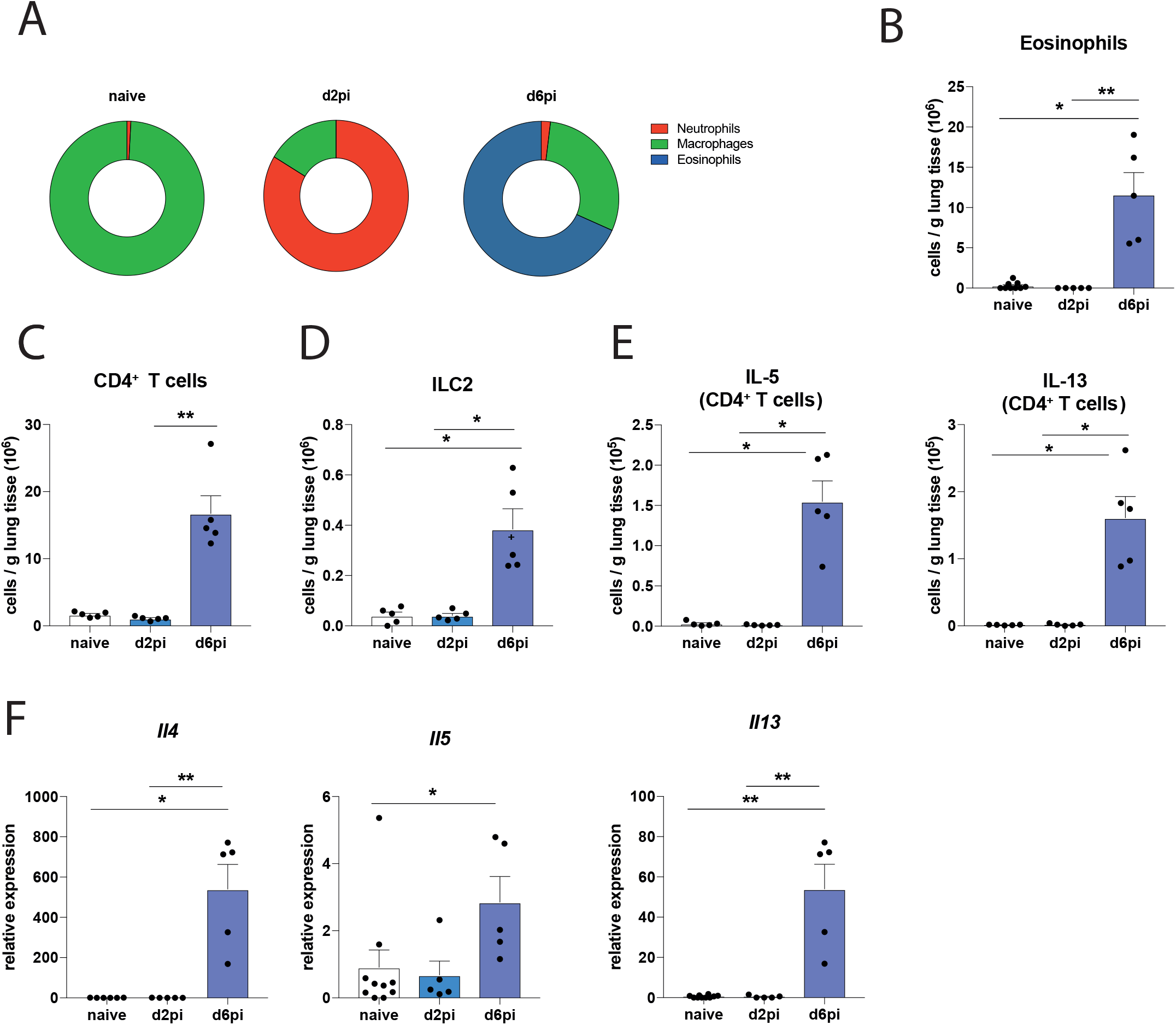
Type-2 immune response induced by *N. brasiliensis* infection. C57BL/6J mice were infected with 500 *N. brasiliensis* L3’s and BAL fluid and whole lung tissue were assessed at d2 and d6 post-infection compared to uninfected (naïve) mice. Cellular composition of CD45^+^ cells in BAL fluid in naïve mice, or mice at d2pi and d6pi (**A**). Absolute number of eosinophils (**B**) CD4^+^ T cells (**C**), ILC2s (**D**) or IL-5^+^ and IL-13^+^ CD4^+^ T cells (**E**) per gram lung tissue. mRNA expression of type-2 cytokines in whole lung following *N. brasiliensis* infection compared to naïve mice (**F**). Data are representative (mean ± s.e.m.) of at least 3 individual experiments. Data was tested for normality using Shapiro-Wilk test and analysed with one-way ANOVA followed by Sidak’s multiple comparisons test for selected groups \**P*<0.05, \*\**P*<0.01.

**Supplement Figure 2:**
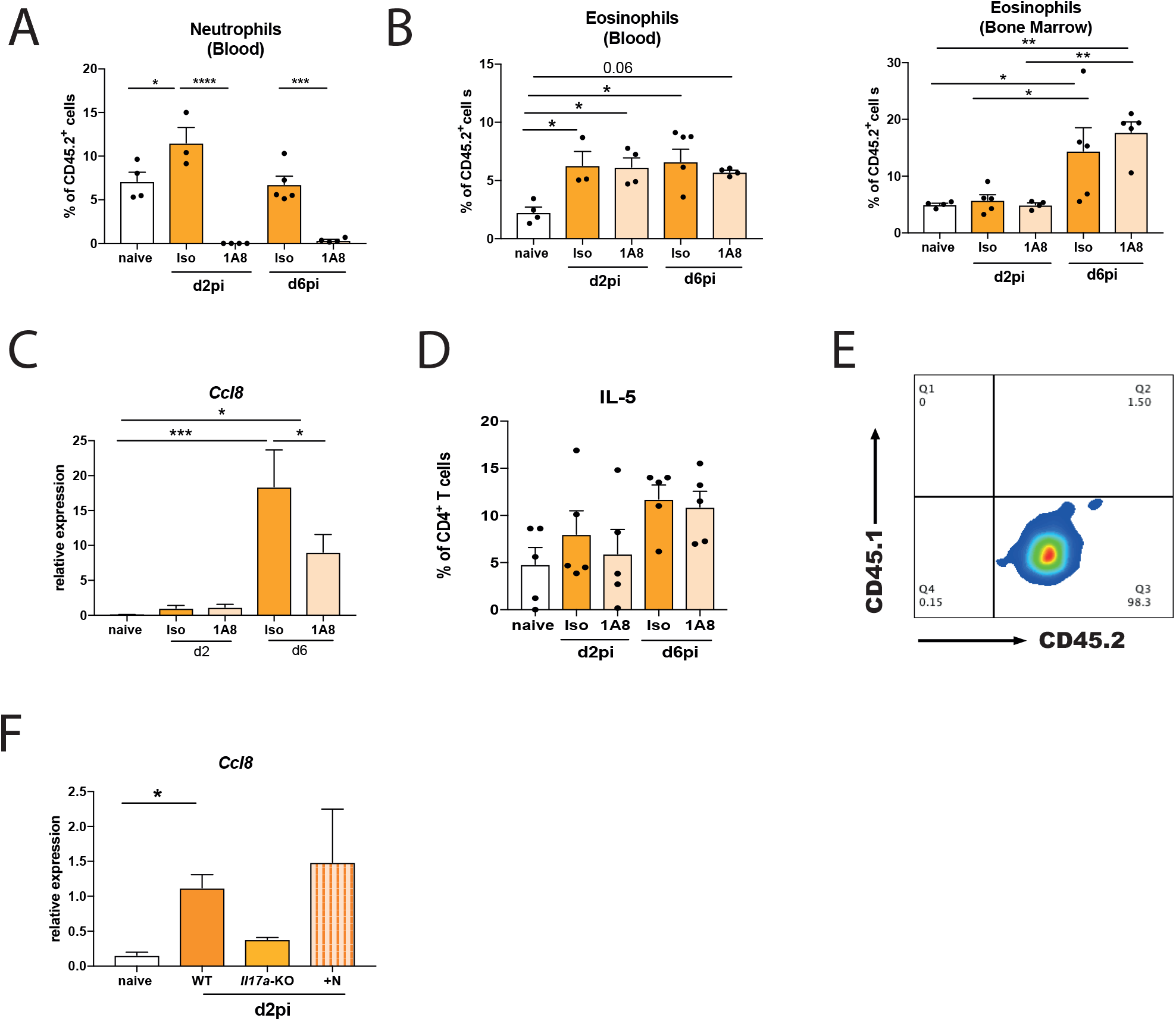
Neutrophils regulate eosinophils in a CCL8-dependent manner. C57BL/6J mice were infected with 500 L3 larvae *N. brasiliensis* and mice were injected with α-Ly6G (1A8) or isotype control on d-1 and d1pi and compared to uninfected, untreated naïve mice. Neutrophils (CD11b^+^ Gr1^+^) in whole blood, showing successful neutrophil depletion (**A**), and eosinophils in blood and bone marrow, expressed as a frequency of CD45.2^+^ cells, in infected mice depleted of neutrophils and isotype controls compared to naïve mice (**B**). Relative mRNA expression of *Ccl8* in whole lung in C57Bl/6J mice after neutrophil depletion and in isotype control compared to naïve mice (**C**). Frequency of IL-5^+^ CD4^+^ T cells in the lung of infected WT mice depleted of neutrophils and WT isotype controls compared to naïve mice (**D**). Representative flow plot showing Ly6G^+^ CD11b^+^ neutrophils in *Il17a*-KO mice 24h after CD45.1^+^ neutrophil transfer into host CD45.2^+^ mice at day 2 post-infection (**E**). Relative mRNA expression of *Ccl8* in whole lung in infected C57Bl/6J mice and *Il17a*-KO mice after neutrophil transfer compared to naïve WT mice (**F**). Data are expressed as mean ± s.e.m. and are representative of at least 3 individual experiments (**A, B, D**) or pooled data from 2 experiments (**C, F**). Data were tested for normality using Shapiro-Wilk test and analysed using a one-way ANOVA followed by Sidak’s multiple comparisons test for selected groups. \**P*<0.05, \*\**P*<0.01, \*\*\**P*<0.001, \*\*\*\**P*<0.0001.

**Supplement Figure 3:**
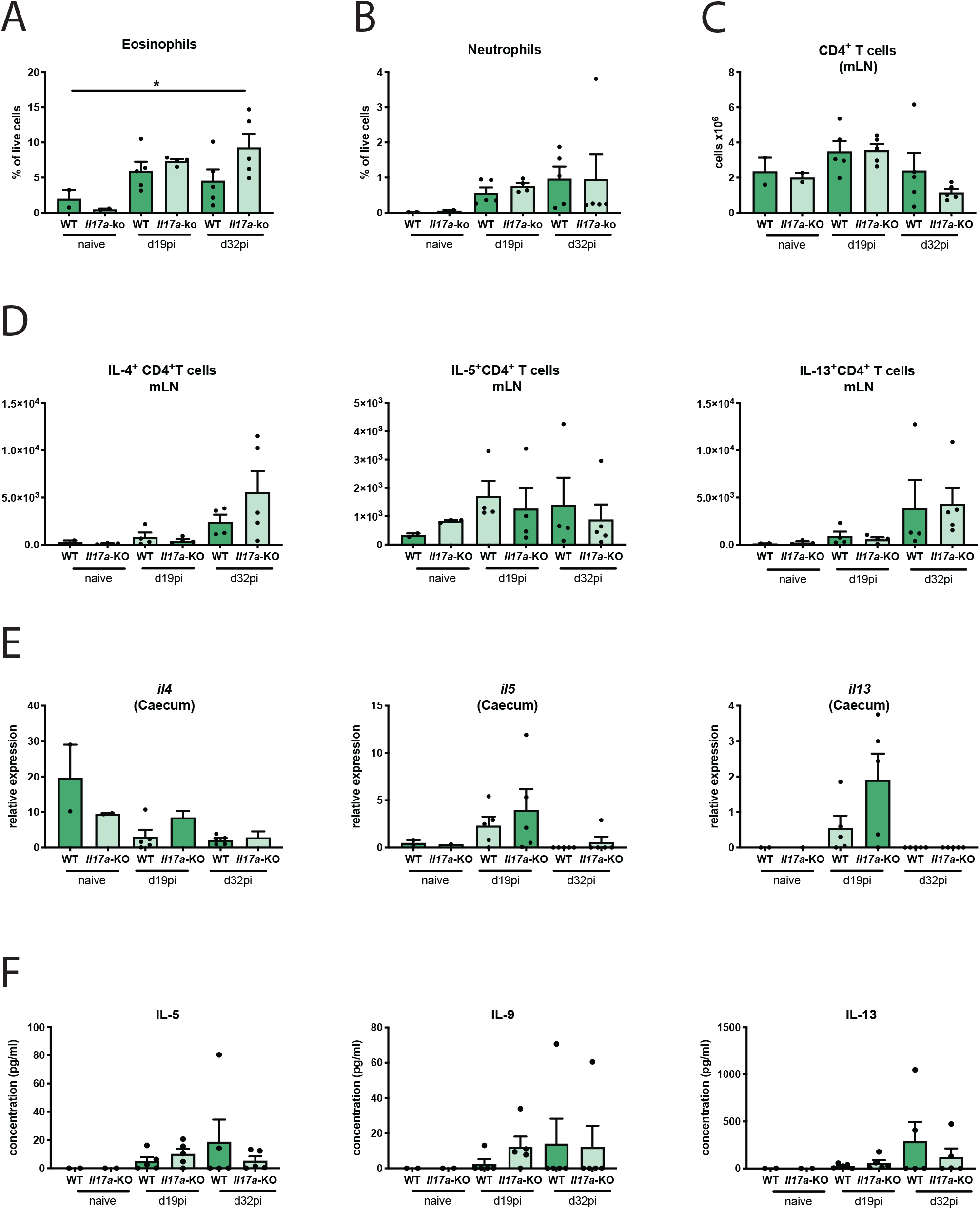
IL-17A is not needed to an induce a protective immune response at the site of infection with *T. muris.* C57Bl/6J and *Il17a*-KO mice were infected with a high dose of *T. muris* (200 larvae) and immune parameters were investigated at d19 and d32pi and compared to uninfected (naïve) C57Bl/6J and *Il17a*-KO mice. Frequencies of eosinophils (**A**) and neutrophils (**B**) in the caecum of WT and *Il17a-KO* mice following infection with *T. muris* compared to naïve mice. Absolute number of CD4^+^ T cells (**C**) and IL-4^+^, IL-5^+^ and IL-13^+^ CD4^+^ T cells in the mLN of infected versus uninfected naïve mice (**D**). Relative mRNA expression of *Il4*, *Il5* and *IL-13* in whole caecum in WT and *Il17a-KO* mice (expression relative to *actb* (β-actin)) (**E**). Protein levels of type-2 cytokines IL-5, IL-9 and IL-13 in mLN recall assays after a 72h restimulation with *T. muris* E/S products (**F**). Data are expressed as mean ± s.e.m. and are representative of at 2 individual experiments. Data was tested for normality using Shapiro-Wilk test and analysed with a one-way ANOVA followed by Sidak’s multiple comparisons test for selected groups; \**P*<0.05.

## REFERENCES

1. Papotto, P. H., Ribot, J. C. & Silva-Santos, B. IL-17 + γδ T cells as kick-starters of inflammation. Nat. Immunol. 18, 604–611 (2017).

2. Iwakura, Y., Ishigame, H., Saijo, S. & Nakae, S. Functional Specialization of Interleukin-17 Family Members. Immunity 34, 149–162 (2011).

3. Allen, J. E. & Maizels, R. M. Diversity and dialogue in immunity to helminths. Nat. Rev. Immunol. 11, 375–388 (2011).

4. Mbow, M. et al. T-helper 17 cells are associated with pathology in human schistosomiasis. J. Infect. Dis. 207, 186–195 (2013).

5. Rutitzky, L. I. & Stadecker, M. J. Exacerbated egg-induced immunopathology in murine Schistosoma mansoni infection is primarily mediated by IL-17 and restrained by IFN-γ. Eur. J. Immunol. 41, 2677–2687 (2011).

6. Katawa, G. et al. Hyperreactive Onchocerciasis is Characterized by a Combination of Th17-Th2 Immune Responses and Reduced Regulatory T Cells. PLoS Negl. Trop. Dis. 9, e3414 (2015).

7. Choy, D. F. et al. TH2 and TH17 inflammatory pathways are reciprocally regulated in asthma. Sci. Transl. Med. 7, 301ra129 (2015).

8. Sutherland, T. E. et al. Chitinase-like proteins promote IL-17-mediated neutrophilia in a tradeoff between nematode killing and host damage. Nat. Immunol. 15, 1116–25 (2014).

9. Molofsky, A. B. et al. InterleuKin-33 And Interferon-Γ Counter-Regulate Group 2 Innate Lymphoid Cell Activation During Immune Perturbation. Immunity 43, 161–174 (2015).

10. Newcomb, D. C. et al. A Functional IL-13 Receptor Is Expressed on Polarized Murine CD4 + Th17 Cells and IL-13 Signaling Attenuates Th17 Cytokine Production. J. Immunol. 182, 5317–5321 (2009).

11. Chen, F. et al. Neutrophils prime a long-lived effector macrophage phenotype that mediates accelerated helminth expulsion. Nat. Immunol. 15, 938–946 (2014).

12. Nakajima, S. et al. IL-17A as an inducer for Th2 immune responses in murine atopic dermatitis models. J. Invest. Dermatol. 134, 2122–2130 (2014).

13. Nakae, S. et al. Antigen-specific T cell sensitization is impaired in Il-17-deficient mice, causing suppression of allergic cellular and humoral responses. Immunity 17, 375–387 (2002).

14. Chenuet, P. et al. Neutralization of either IL-17A or IL-17F is sufficient to inhibit house dust mite induced allergic asthma in mice. Clin. Sci. 131, 2533–2548 (2017).

15. Campbell, L. et al. ILC2s mediate systemic innate protection by priming mucus production at distal mucosal sites. J. Exp. Med. 216, 2714–2723 (2019).

16. Minutti, C. M. et al. Epidermal Growth Factor Receptor Expression Licenses Type-2 Helper T Cells to Function in a T Cell Receptor-Independent Fashion. Immunity 47, 710–722.e6 (2017).

17. Redpath, S. A. et al. ICOS controls Foxp3+ regulatory T-cell expansion, maintenance and IL-10 production during helminth infection. Eur. J. Immunol. 43, 705–715 (2013).

18. Terrazas, L. I., Montero, D., Terrazas, C. A., Reyes, J. L. & Rodríguez-Sosa, M. Role of the programmed Death-1 pathway in the suppressive activity of alternatively activated macrophages in experimental cysticercosis. Int. J. Parasitol. 35, 1349–1358 (2005).

19. Nembrini, C., Marsland, B. J. & Kopf, M. IL-17-producing T cells in lung immunity and inflammation. J. Allergy Clin. Immunol. 123, 986–994 (2009).

20. Porzionato, A. et al. Fractal analysis of alveolarization in hyperoxia-induced rat models of bronchopulmonary dysplasia. Am. J. Physiol. – Lung Cell. Mol. Physiol. 310, L680–L688 (2016).

21. Chenery, A. L. et al. Inflammasome-Independent Role for NLRP3 in Controlling Innate Antihelminth Immunity and Tissue Repair in the Lung. J. Immunol. 203, 2724–2734 (2019).

22. Selders, G. S., Fetz, A. E., Radic, M. Z. & Bowlin, G. L. An overview of the role of neutrophils in innate immunity, inflammation and host-biomaterial integration. Regen. Biomater. 4, 55–68 (2017).

23. Ardain, A. et al. Group 3 innate lymphoid cells mediate early protective immunity against tuberculosis. Nature 570, 528–532 (2019).

24. Osborne, L. C. et al. Coinfection. Virus-helminth coinfection reveals a microbiota-independent mechanism of immunomodulation. Science (80-.). 345, 578–582 (2014).

25. Ribot, J. C. et al. CD27 is a thymic determinant of the balance between interferon-γ- and interleukin 17-producing γδ T cell subsets. Nat. Immunol. 10, 427–436 (2009).

26. Bancroft, A. J., McKenzie, A. N. & Grencis, R. K. A critical role for IL-13 in resistance to intestinal nematode infection. J. Immunol. 160, 3453–61 (1998).

27. Bancroft, A. J., Else, K. J. & Grencis, R. K. Low-level infection with Trichuris muris significantly affects the polarization of the CD4 response. Eur. J. Immunol. 24, 3113–3118 (1994).

28. Filbey, K. J. et al. Intestinal helminth infection promotes IL-5- and CD4 + T cell-dependent immunity in the lung against migrating parasites. Mucosal Immunol. 12, 352–362 (2019).

29. Lajoie, S. et al. Complement-mediated regulation of the IL-17A axis is a central genetic determinant of the severity of experimental allergic asthma. Nat. Immunol. 11, 928–935 (2010).

30. Díaz, A. & Allen, J. E. Mapping immune response profiles: The emerging scenario from helminth immunology. Eur. J. Immunol. 37, 3319–3326 (2007).

31. Wakashin, H., Hirose, K., Iwamoto, I. & Nakajima, H. Role of IL-23-Th17 cell axis in allergic airway inflammation. In Int. Arch. Allergy Immunol. (2009).doi:10.1159/000211382

32. Wang, X. et al. IL-17 constrains natural killer cell activity by restraining IL-15-driven cell maturation via SOCS3. Proc. Natl. Acad. Sci. U. S. A. 116, 17409–17418 (2019).

33. Moroda, M., Takamoto, M., Iwakura, Y., Nakayama, J. & Aosai, F. Interleukin-17Adeficient mice are highly susceptible to Toxoplasma gondii infection due to excessively induced T. gondii HSP70 and interferon gamma production. Infect. Immun. 85, (2017).

34. Ritter, M. et al. Absence of IL-17A in Litomosoides sigmodontis-infected mice influences worm development and drives elevated filarial-specific IFN-γ. Parasitol. Res. 117, 2665–2675 (2018).

35. Zhang, Y. et al. Lack of IL-17 signaling decreases liver fibrosis in murine schistosomiasis japonica. Int. Immunol. 27, 317–325 (2015).

36. Li, L. et al. IL-17 produced by neutrophils regulates IFN-γ-mediated neutrophil migration in mouse kidney ischemia-reperfusion injury. J. Clin. Invest. 120, 331–342 (2010).

37. Lin, Y. et al. Interleukin-17 Is Required for T Helper 1 Cell Immunity and Host Resistance to the Intracellular Pathogen Francisella tularensis. Immunity 31, 799–810 (2009).

38. Taylor, B. C. et al. TSLP regulates intestinal immunity and inflammation in mouse models of helminth infection and colitis. J. Exp. Med. 206, 655–667 (2009).

39. Costa Carvalho, J. L. et al. The chemokines secretion and the oxidative stress are targets of low-level laser therapy in allergic lung inflammation. J. Biophotonics 9, 1208–1221 (2016).

40. Dulek, D. E. et al. Allergic airway inflammation decreases lung bacterial burden following acute Klebsiella pneumoniae infection in a neutrophil- and CCL8-dependent manner. Infect. Immun. 82, 3723–3739 (2014).

41. Puttur, F. et al. Pulmonary environmental cues drive group 2 innate lymphoid cell dynamics in mice and humans. Sci. Immunol. 4, eaav7638 (2019).

42. Newcomb, D. C. et al. IL-13 Regulates Th17 Secretion of IL-17A in an IL-10– Dependent Manner. J. Immunol. 188, 1027–1035 (2012).

43. Dyken, S. J. Van et al. Chitin activates parallel immune modules that direct distinct inflammatory responses via innate lymphoid type 2 and γδ T cells. Immunity 40, 414–424 (2014).

44. Chen, F. et al. An essential role for T H 2-type responses in limiting acute tissue damage during experimental helminth infection. Nat. Med. 18, 260–266 (2012).

45. Srinivas, S. et al. Cre reporter strains produced by targeted insertion of EYFP and ECFP into the ROSA26 locus. BMC Dev. Biol. 1, 1–8 (2001).

46. Hirota, K. et al. Fate mapping of IL-17-producing T cells in inflammatory responses. Nat. Immunol. 12, 255–263 (2011).

47. Lawrence, R. A., Gray, C. A., Osborne, J. & Maizels, R. M. Nippostrongylus brasiliensis: Cytokine responses and nematode expulsion in normal and IL-4 deficient mice. Exp. Parasitol. 84, 65–73 (1996).

48. Holland, M. J., Harcus, Y. M., Riches, P. L. & Maizels, R. M. Proteins secreted by the parasitic nematode Nippostrongylus brasiliensis act as adjuvants for Th2 responses. Eur. J. Immunol. 30, 1977–1987 (2000).

49. Swamydas, M., Luo, Y., Dorf, M. E. & Lionakis, M. S. Isolation of mouse neutrophils. Curr. Protoc. Immunol. 2015, 3.20.1–3.20.15 (2015).

50. Hayes, K. S. et al. Chronic Trichuris muris infection causes neoplastic change in the intestine and exacerbates tumour formation in APC min/+ mice. PLoS Negl. Trop. Dis. 11, (2017).

51. Ritchie, M. E. et al. Limma powers differential expression analyses for RNA-sequencing and microarray studies. Nucleic Acids Res. 43, e47 (2015).

